# CD22L Conjugation to Insulin Attenuates Insulin-Specific B cell Activation

**DOI:** 10.1101/2023.06.13.544802

**Authors:** Kyle D. Apley, Amber S. Griffith, Grant M. Downes, Patrick Ross, Mark P. Farrell, Peggy Kendall, Cory J. Berkland

**Affiliations:** Department of Pharmaceutical Chemistry, University of Kansas, Lawrence, Kansas 66047, United States; Department of Medicine, Division of Allergy and Immunology, Washington University School of Medicine, St. Louis, Missouri, 63110, United States; Bioengineering Program, University of Kansas, Lawrence, Kansas 66045, United States; Department of Medicinal Chemistry, University of Kansas, Lawrence, Kansas 66047, United States; Department of Chemical and Petroleum Engineering, University of Kansas, Lawrence, Kansas 66045, United States; Department of Biomedical Engineering, Washington University, St. Louis, Missouri 63130, United States; Department of Chemistry, Washington University, St. Louis, Missouri 63130, United States

**Keywords:** Insulin, CD22, Type 1 Diabetes, B cell, tolerance, antigen

## Abstract

Pancreatic islet-reactive B lymphocytes promote Type 1 diabetes (T1D) by presenting antigen to islet-destructive T cells. Teplizumab, an anti-CD3 monoclonal, delays T1D onset in patients at risk, but additional therapies are needed to prevent disease entirely. Therefore, bifunctional molecules were designed to selectively inhibit T1D-promoting anti-insulin B cells by conjugating a ligand for the B cell inhibitory receptor CD22 (i.e., CD22L) to insulin, which permit these molecules to concomitantly bind to anti-insulin B cell receptors (BCRs) and CD22. Two prototypes were synthesized: 2:2 insulin-CD22L conjugate on a 4-arm PEG backbone, and 1:1 insulin-CD22L direct conjugate. Transgenic mice (125TgSD) expressing anti-insulin BCRs provided cells for in vitro testing. Cells were cultured with constructs for three days then assessed by flow cytometry. Duplicate wells with anti-CD40 simulated T cell help. Surprisingly, a 2-insulin 4-arm PEG control caused robust proliferation and activation-induced CD86 upregulation. Anti-CD40 further boosted these effects. This was unexpected, as soluble insulin alone has no effect, and may indicate that BCR-crosslinking occurs when antigens are tethered by the PEG backbone. Addition of CD22L via the 2:2 insulin-CD22L conjugate restored B cell properties to that of controls without additional beneficial effect. In contrast, the 1:1 insulin-CD22L direct conjugate significantly reduced anti-insulin B cell proliferation in the presence of anti-CD40. CD22L alone had no effect, and the constructs did not affect WT B cells. Thus, high valency constructs activate anti-insulin B cells, while low-valency antigen-CD22L conjugates co-ligate BCR and CD22, reducing B cell activation in response to simulated T cell help and reducing pathogenic B cell numbers without harming normal cells. Thus, the insulin-CD22L direct conjugate is a promising candidate for preclinical trials to prevent T1D without inducing immunodeficiency

## Introduction

Autoreactive B cells contribute to aberrant immune responses in autoimmune disease through autoantibody production, autoantigen presentation to T cells, and inflammatory cytokine production.^1, 2^ One therapeutic strategy to restore B cell tolerance in autoimmunity is to block B cell activation by co-localizing the B cell receptor (BCR) with intracellular immunoreceptor tyrosine-based inhibitory motif (ITIM)-bearing receptors using antigen-ligand or anti-BCR antibody-ligand conjugates.^3–15^ ITIM-bearing receptors expressed on B cells investigated in this approach include the siglecs CD33 and CD22 and the antibody-Fc receptor FcγRIIB. Notably, targeting of CD22 may contribute to selective B cell immunomodulation due to its sole expression on B cells.^16, 17^ Co-localization of CD22 with the BCR complex results in phosphorylation of the CD22 ITIM by the Src-family protein tyrosine kinase LYN followed by activation of the protein tyrosine phosphatase SHP-1.^18, 19^ SHP-1 then negatively regulates B cell activation by suppressing downstream effectors including MAP kinases and phospholipase C responsible for intracellular calcium release.^20^

Prior tolerization studies with BCR – CD22 co-localization focused on disease models in which antibodies are key effectors of disease including rheumatoid arthritis and alloantibody-mediated hemophilia, and these studies primarily reported *in vivo* antibody titer data.^5–10^ In type 1 diabetes (T1D), however, autoantibodies do not have such a clear role in disease although their presence indicates risk of disease development. On the other hand, effector anti-islet antigen B cells are indeed pathogenic, propagating autoimmunity via autoantigen presentation and inflammatory cytokine excretion.^21–31^ In addition, these prior B cell tolerization studies relied on liposome and large polymer scaffolds bearing hundreds of copies of antigen and CD22 ligand (CD22L), and it is unknown if low valency antigen-CD22L conjugates have the potential to tolerize B cells in a similar manner.

Currently, there is little data on antigen-CD22L conjugates’ ability to block B-cell activation in relation to T1D. For these studies, we developed therapeutic bifunctional molecules designed to target anti-insulin B cells that promote T1D. These bifunctional molecules use insulin autoantigen to bind the BCR and are conjugated with murine CD22L (mCD22L), developed by Paulson and co-workers to bind murine CD22 with micromolar affinity.^32^ Transgenic anti-insulin (125Tg) B cells were used to study effects of low valency conjugates with a discrete number of insulin molecules and mCD22Ls (1:1, 2:2). The 1:1 insulin-mCD22L conjugate was synthesized by direct conjugation of an azide-functionalized mCD22L to insulin(B29Lys)-alkyne, and the 2:2 insulin-mCD22L conjugate utilized a 20 kDa 4-arm PEG scaffold. Following characterization, the conjugates were incubated with 125Tg splenocytes with and without anti-CD40 to investigate the effect of the conjugates on B cell survival, proliferation and CD86 expression. The data demonstrate the ability of CD22L and antigen conjugation to attenuate anti-insulin B cell activation *in vitro*.

## Methods

### Insulin-azide, insulin-alkyne, and mCD22L-azide synthesis

Insulin-azide and insulin-alkyne were prepared from recombinant human insulin (EMD Millipore) by selective acylation of the B29Lys residue as previously reported.^33, 34^ The synthesis of mCD22L-azide is detailed in the supporting information.

### 4-arm PEG-insulin synthesis

To a solution of 40 mg of 4-arm PEG-alkyne 20kDa (MW = 20000, 2 μmoles, Advanced Biochemicals) in 3.4 mL of 50 mM phosphate pH 7.4 buffer, 24.3 mg of insulin-azide (MW = 6080.92, 4 μmoles) in 1 mL of DMSO was added followed by 1 mg of CuSO_4_ •7H_2_O (MW = 249.69, 4 μmoles) and 8.7 mg of THPTA (MW = 434.50, 20 μmoles) dissolved in 400 μL of water. Lastly, 15.85 mg of sodium ascorbate (MW = 198.11, 80 μmoles) dissolved in 200 μL of water was added to the reaction. The reaction was stirred at RT for 1 hour. The resulting 4-arm PEG-insulin products with 1, 2, 3, or 4 insulin per PEG were isolated by preparative reverse-phase HPLC (Prep-LC) using a Waters XBridge Prep C18 5 μm OBD 19 x 250 mm column at a flow rate of 14 ml/min with a 10-minute gradient from 36-42%B (solvent A: water with 0.05% trifluoroacetic acid (TFA), solvent B: acetonitrile with 0.05% TFA). Desired fractions were concentrated by rotary evaporation under reduced pressure, frozen, then lyophilized to give a white powder. Yields of 7.1, 10.1, 11.7, and 8.2 mgs were attained. Product identity was confirmed by matrix-assisted laser desorption/ionization time-of-flight (MALDI-TOF) mass spectrometry using a HABA matrix. Expected: 26,000, 32,000, 38,000, 44,000 Da; found: 26,081, 32,332, 38,096, 44,130 m/z [M+H]^+^.

### 4-arm PEG-insulin(2)/mCD22L(2) synthesis

To a solution of 4.3 mg of 4-arm PEG-insulin(2) (MW = 32,000, 0.134 μmoles) dissolved in 1 mL of 50 mM phosphate pH 7.4 buffer with 250 μL of DMSO, 0.54 mg of mCD22L-azide (MW = 999.97, 0.54 μmoles) in 81 μL of buffer was added followed by 67 μg of CuSO4 •7H2O (MW = 249.69, 0.268 μmoles) and 587 μg of THPTA (MW = 434.50, 1.34 μmoles) dissolved in 27 μL of water. Lastly, 1.07 mg of sodium ascorbate (MW = 198.11, 5.36 μmoles) dissolved in 13.5 μL of water was added to the reaction. The reaction was stirred at RT for 2 hours. The desired product was isolated by Prep-LC using a Waters XBridge Prep C18 5 μm OBD 19 x 250 mm column at a flow rate of 14 ml/min with a 10-minute gradient from 35-45%B (solvent A: water with 0.05% trifluoroacetic acid (TFA), solvent B: acetonitrile with 0.05% TFA). Desired fractions were concentrated by rotary evaporation under reduced pressure, frozen, then lyophilized to give a white powder. Yield: 2.45 mgs (55%). Product identity was confirmed by matrix-assisted laser desorption/ionization time-of-flight (MALDI-TOF) mass spectrometry using a DHAP matrix. Expected: 34,000 Da. Found: 34,140 m/z [M+H]+.

### Insulin-mCD22L synthesis

To a solution of 4 mg of insulin-alkyne (MW = 5916.77, 0.68 μmoles) dissolved in 340 μL of DMSO and 1.2 mL of 50 mM phosphate pH 7.4 buffer, 0.81 mg of mCD22L-azide (MW = 999.97, 0.81 μmoles) in 81 μL of buffer was added followed by 170 μg of CuSO4 •7H2O (MW = 249.69, 0.68 μmoles) and 1.47 mg of THPTA (MW = 434.50, 3.4 μmoles) dissolved in 68 μL of water. Lastly, 2.68 mg of sodium ascorbate (MW = 198.11, 13.6 μmoles) dissolved in 34 μL of water was added to the reaction. The reaction was stirred at RT for 1 hour. The desired product was isolated by Prep-LC using a Waters XBridge Prep C18 5 μm OBD 19 x 250 mm column at a flow rate of 14 ml/min with a 10-minute gradient from 20-40%B (solvent A: water with 0.05% trifluoroacetic acid (TFA), solvent B: acetonitrile with 0.05% TFA). Desired fractions were concentrated by rotary evaporation under reduced pressure, frozen, then lyophilized to give a white powder. Yield: 3.54 mgs (76%). Product identity was confirmed by matrix-assisted laser desorption/ionization time-of-flight (MALDI-TOF) mass spectrometry using a DHAP matrix. Expected: 6917.7 Da. Found: 6918.06 m/z [M+H]+.

### RP-HPLC Analysis

Lyophilized Prep-LC fractions dissolved in DMSO/water at 0.5 mg/ml were injected onto a Waters XBridge BEH C18 2.5 μm 130 Å 4.6 x 150 mm column equilibrated at 50 °C using a 10 minute 30-50%B gradient (solvent A: water with 0.05% trifluoroacetic acid (TFA), solvent B: acetonitrile with 0.05% TFA). Detection was performed at 214 and 280 nm.

### B Cell Proliferation and Flow Cytometry

Splenocytes from 125Tg B6 and WT B6 control mice were stained with CellTrace Violet and cultured for 3 days^35^ in complete RPMI alone or stimulated with 5 ug/mL anti-mouse anti-CD40 (BD Pharmingen clone HM40-3) in the presence or absence of conjugates described above at 150 nM concentrations. The cells were then analyzed by flow cytometry using fluorochrome-conjugated antibodies against B220 (RA3-6B2, BUV395), CD86 (GL-1 PE), and CD22 (Cy34.1 FITC) from BD Biosciences. LiveDead Blue, CellTrace Violet and were obtained from Invitrogen. APC conjugated IgMa was obtained from Biolegend. Samples were collected on an Aurora Flow Cytotmeter (Cytek Biosciences), and data was analyzed using Flowjo software (Tree Star) and GraphPad Prism.

## Results

### Insulin-mCD22L Conjugate Synthesis

Insulin-mCD22L was prepared by copper-catalyzed azide-alkyne cycloaddition (CuAAC) involving insulin-alkyne and mCD22L-azide (**Figure 1**). The desired product was isolated by preparatory RP-HPLC and product identity was confirmed by MALDI-TOF mass spectrometry. The 4-arm PEG-insulin(2)/mCD22L(2) was prepared in a two-step process (**Figure 2**). First, insulin-azide was conjugated to 4-arm PEG-alkyne by CuAAC yielding a mixture of 4-arm PEG-insulin products with 1 to 4 insulin molecules attached per PEG backbone (**Figure 3A, B**). The individual species were isolated using a preparatory reverse-phase HPLC (Prep-LC) method with a shallow gradient. Each fraction was analyzed by HPLC, and the four major products were resolved using a 2.5 µm C18 column at 50 °C. The purity of each species was greater than 90% following Prep-LC purification. The individual 4-arm PEG-insulin species were identified by MALDI mass spectrometry (**Figure 3C**). Due to the inherent MW distribution of the 4-arm PEG-alkyne (20 kDa) backbone, the 4-arm PEG-insulin species also exist with a similar distribution of MWs. Even so, the mode molecular weight for each 4-arm PEG-insulin species closely matched the theoretical MWs (**Figure 3A**).

**Figure 1:**
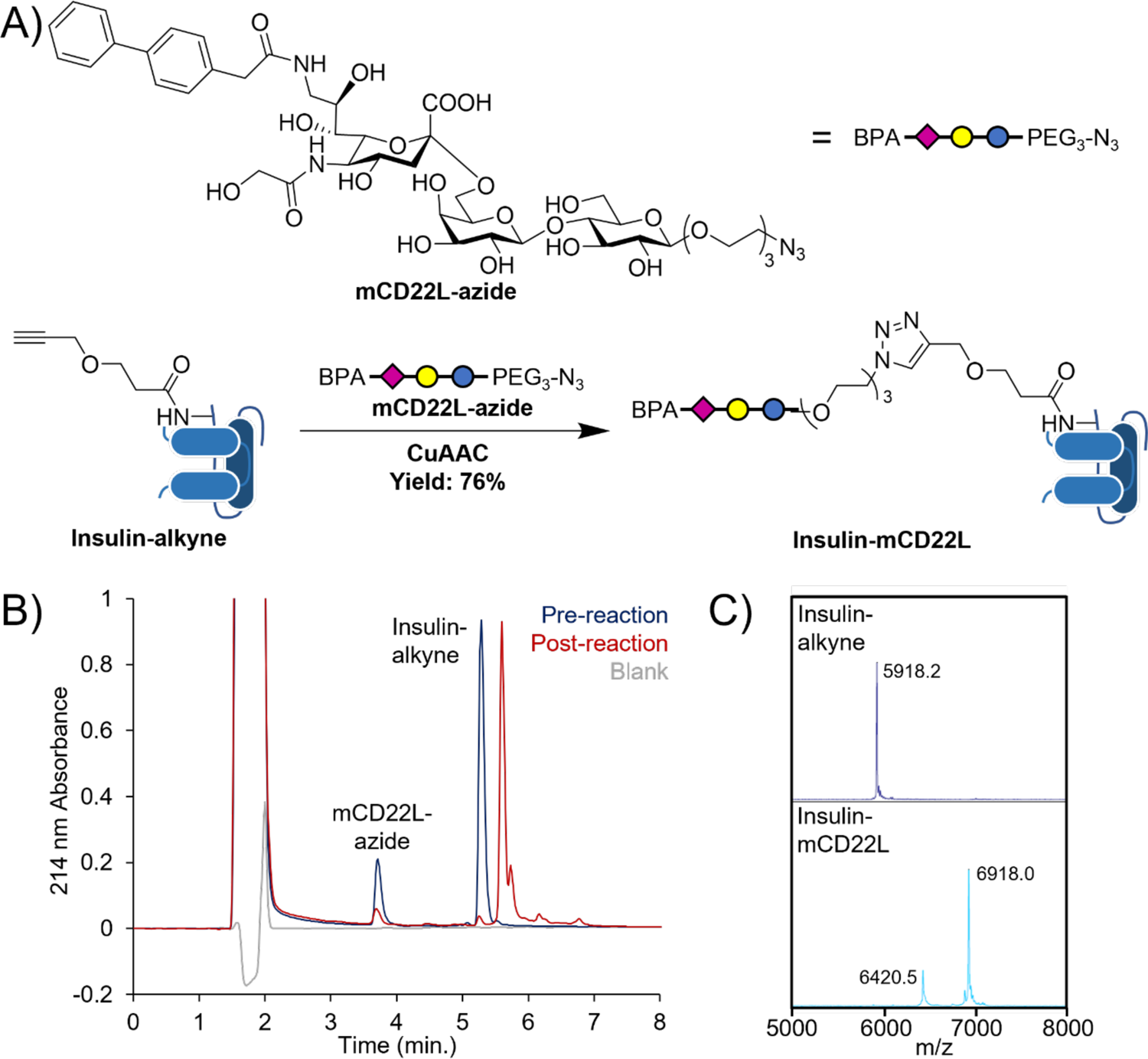
Synthesis of insulin-mCD22L; a direct conjugate between insulin-alkyne and mCD22L-azide. A) Reaction scheme. B) HPLC analysis of reactants and reaction products. C) MALDI-TOF-MS analysis of insulin-alkyne and isolated insulin-mCD22L.

**Figure 2:**
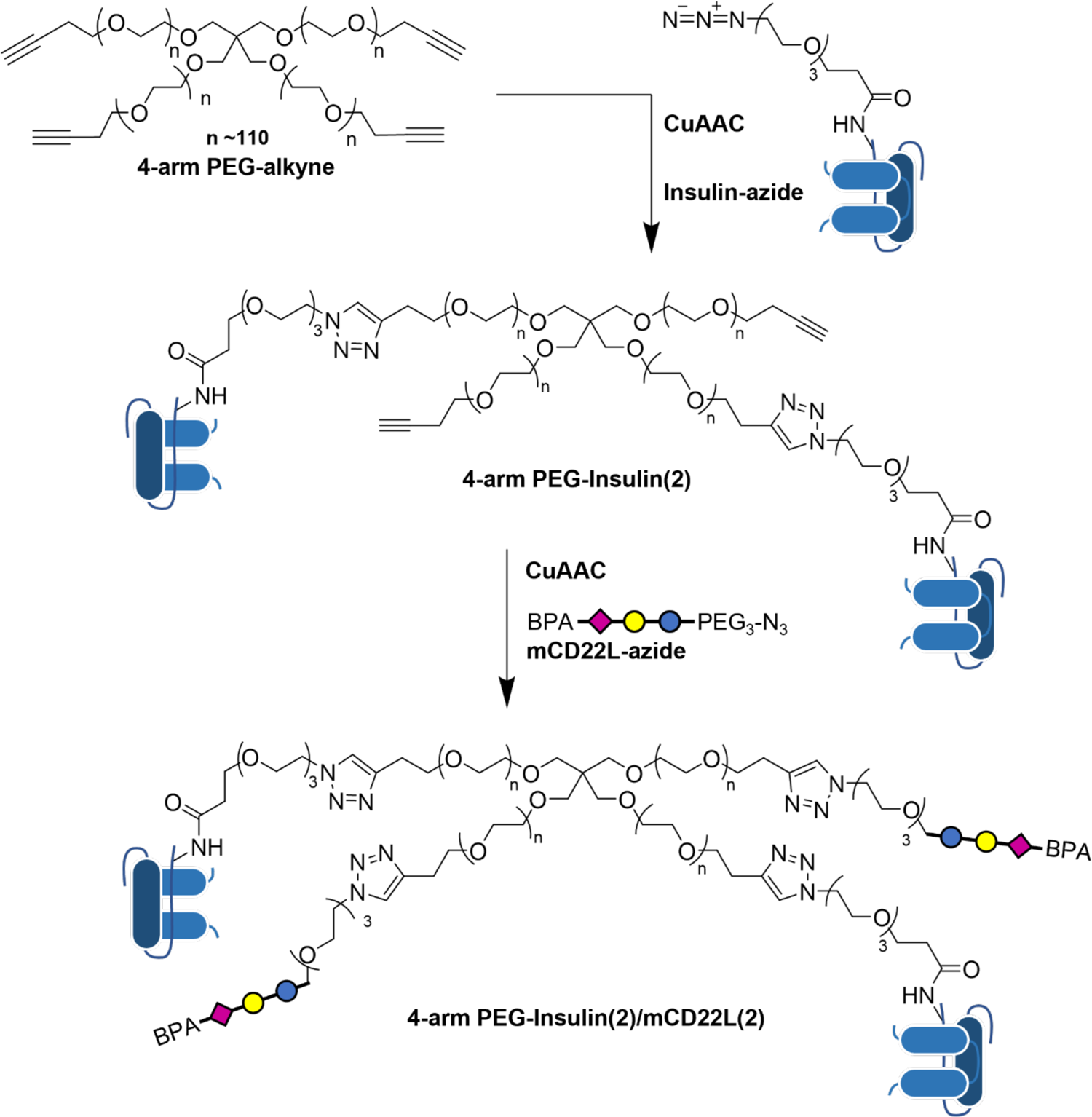
Reaction scheme of 4-arm PEG-insulin(2)/mCD22L(2). First, insulin-azide is conjugated with 4-arm PEG-alkyne by copper-catalyzed azide-alkyne cycloaddition (CuAAC) to yield a mixture of 4-arm PEG-insulin(1-4) products, including 4-arm PEG-insulin(2). Second, isolated 4-arm PEG-insulin(2) is conjugated to mCD22L-azide by CuAAC to yield 4-arm PEG-insulin(2)/mCD22L(2).

**Figure 3:**
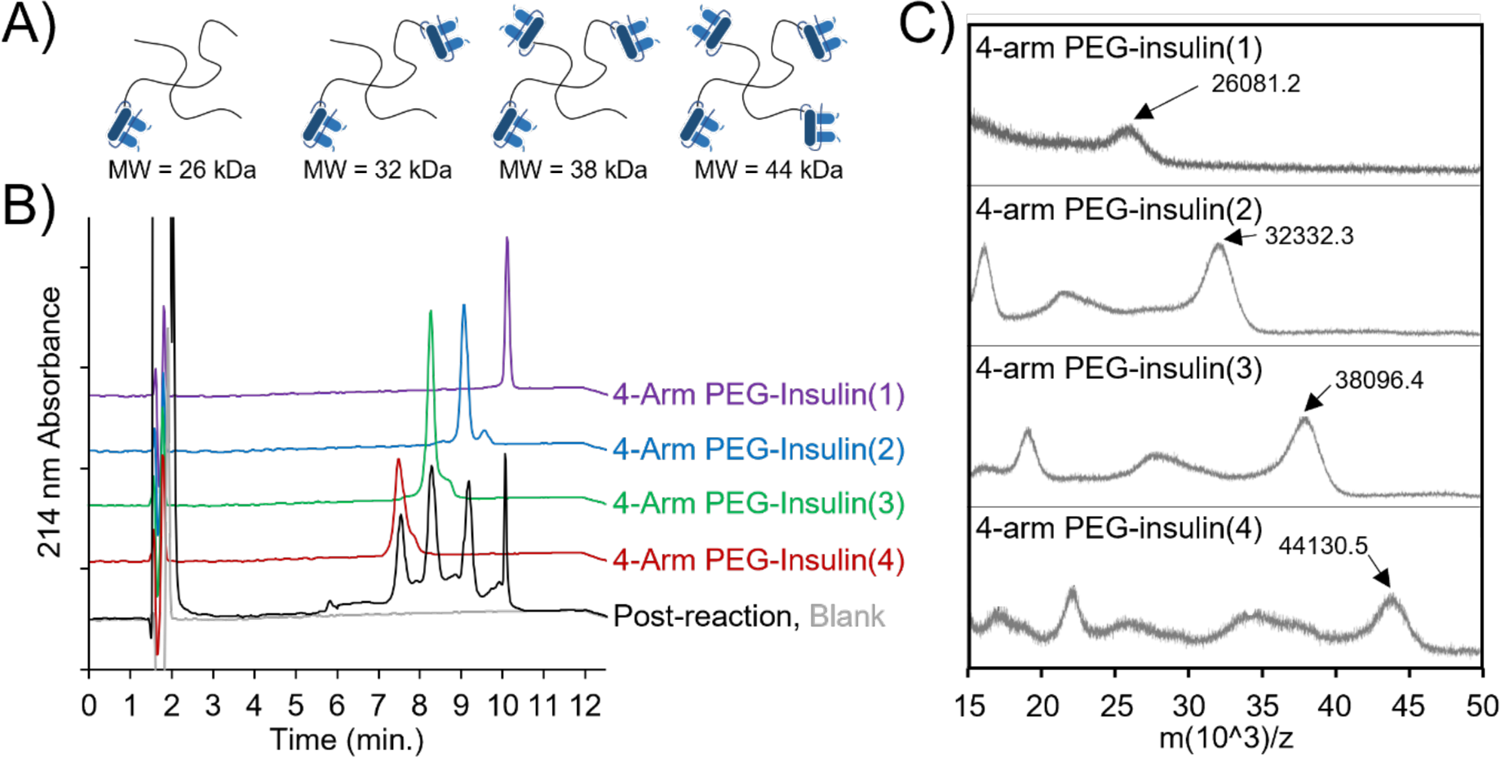
Analysis of 4-arm PEG-insulin products. A) Representation and molecular weights (MW) of the reaction products formed in the CuAAC reaction between 4-arm PEG-alkyne and insulin-azide. B) HPLC analysis of four product fractions following preparatory RP-HPLC purification of 4-arm PEG-insulin. The number in parenthesis indicates the number of insulin molecules conjugated to the 4-arm PEG. C) MALDI-MS analysis of four product fractions following preparatory RP-HPLC purification of 4-arm PEG-insulin. The mode m/z ratio of the major peak is indicated.

Second, the 4-arm PEG-insulin(2) was reacted with mCD22L-azide by CuAAC to prepare a 4-arm PEG-insulin(2)/mCD22L(2) conjugate (Figure 2). Comparison of HPLC traces for pre- and post-reaction samples show a distinct shift in retention time for the product (Figure 4A). MALDI mass spectrometry analysis of the product identified a 2 kDa increase in mode MW, matching the expected 1 kDa increase per mCD22L-azide conjugated (Figure 4B). Proton NMR characterization was performed for 4-arm Peg-Insulin(2)/mCD22L(2) (Figure 4C). Focusing on the aromatic region, peaks from the aromatic amino acids of insulin (6.5-7.5 ppm) are present along with peaks from the bisphenyl-aryl group on the mCD22L (7.5-7.8 ppm).

**Figure 4:**
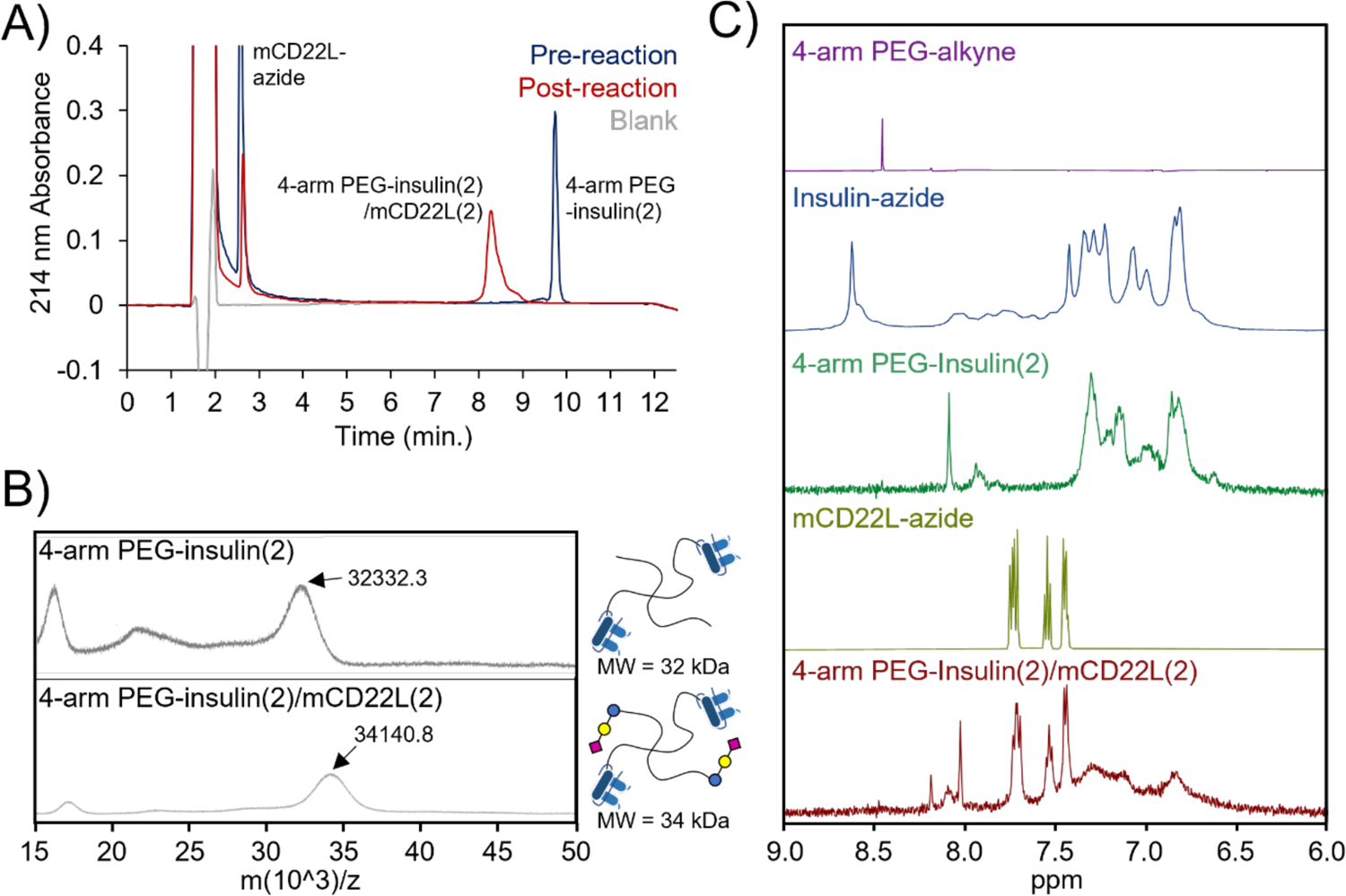
Analysis of 4-arm PEG-insulin/mCD22L products. A) HPLC analysis of the CuAAC reaction between 4-arm PEG-insulin(2) with mCD22L-azide before and 2 hours after the addition of sodium ascorbate. Two equivalents of mCD22L-azide per alkyne were used. B) MALDI-MS analysis of the 4-arm PEG-insulin(2)/mCD22L(2) product following purification by RP-HPLC. The mode m/z ratio of the major peak is indicated. A representation and molecular weights (MW) of the starting material and product are shown. C) Proton NMR analysis of 4-arm PEG-insulin(2)/mCD22L(2) and precursors. The aromatic region of the spectra is shown due to the distinctive peaks from both the aromatic sidechains of insulin-azide and the bisphenyl group on mCD22L-azide.

Additionally, a peak corresponding to the sole proton of the 1,2,3-N triazole formed by CuAAC is present (8.0 ppm). Integration of the mCD22L aromatic peak area and comparison to the 1,2,3-N triazole proton peak and PEG-backbone proton peak support the presence of two mCD22Ls per backbone (**Table S1**). The broadening of the insulin aromatic amino acid peaks made quantitative assessment of the insulins per backbone infeasible for the 4-arm PEG-insulin(2)/mCD22L(2) species, but integration of the insulin aromatic amino acid peak area in the 4-arm PEG-insulin(2) species and comparison to the 1,2,3-N triazole proton peak and PEG-backbone proton peak support the presence of two insulins per backbone, along with the MALDI data (Figure 4C). Together, the HPLC, MS, and NMR data confirm the 4-arm PEG-insulin(2)/mCD22L(2) product to have a discrete number of insulin molecules and mCD22Ls.

### Effect of Insulin-mCD22L Conjugates on B cells

In 125Tg mice, the transgene is expressed by over 95% of B cells, providing a large population of anti-insulin B cells for study.^30^ Splenocytes from 125Tg B6 mice were pre-labeled with the proliferation dye Cell Trace Violet then incubated with insulin constructs at 150 nM concentration.^35^ Following 72 hours of incubation, flow cytometry was used to measure construct effects on B cell survival, proliferation and activation. Surprisingly, the 4-arm PEG-insulin(2) control construct had a strong activating effect on anti-insulin B cells (Figure 5A), resulting in a significantly higher proportion remaining in wells compared to those in control wells (11.99% ± 8.24% SD; media only 3.45% ± 2.53% SD; mCD22L 3.99% ± 2.34% SD).

**Figure 5:**
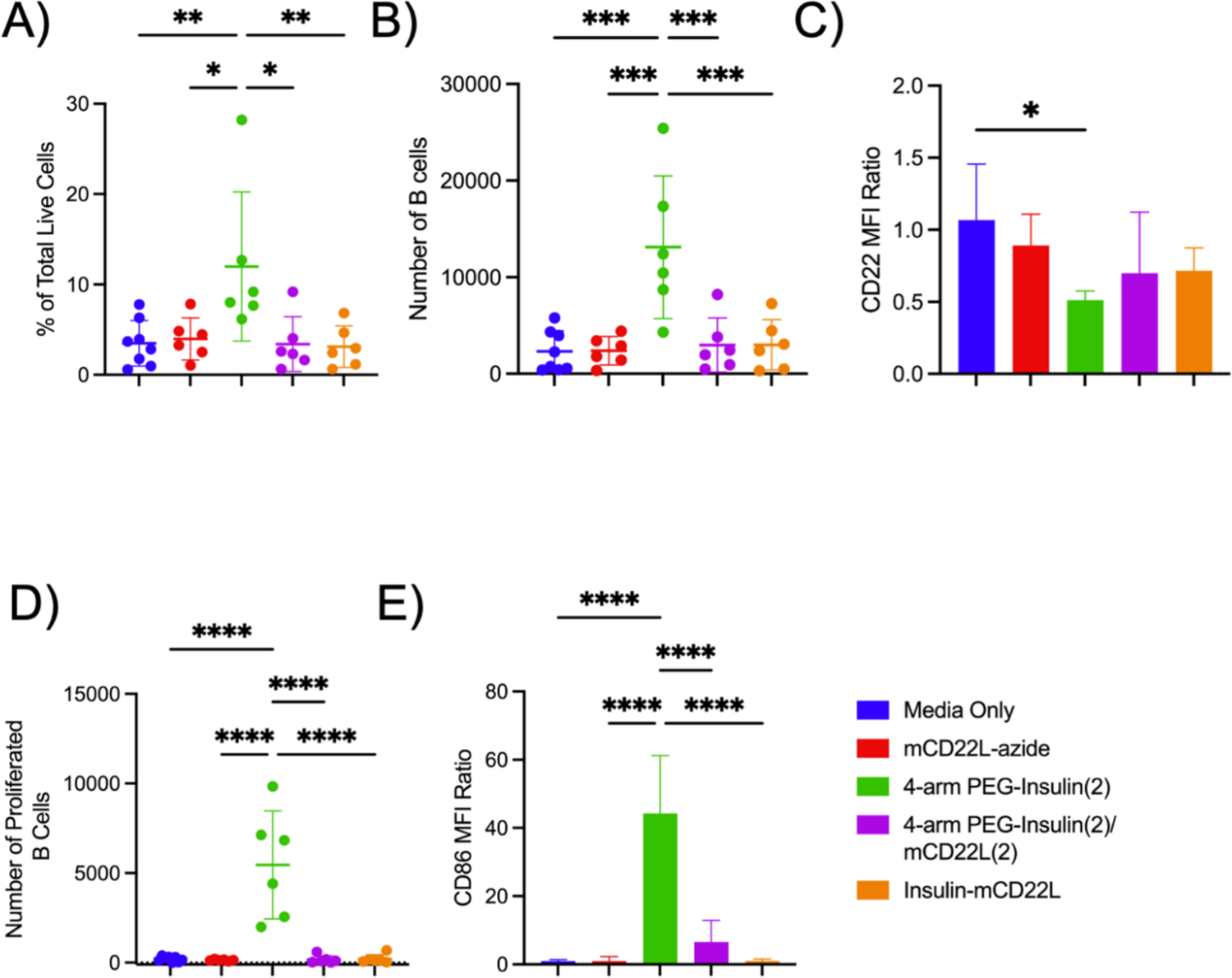
Anti-insulin B cell responses to the incubation of insulin and mCD22L conjugates in 125Tg B6 mouse splenocytes after 72 hours. A) The percentage of total live cells that are B cells. B) Total live B cell number. C) CD22 expression levels on B cells as a ratio over untreated anti-insulin B cell CD22 expression. D) Number of proliferating B cells determined by dye-dilution assay. E) CD86 expression levels on B cells as a ratio over untreated anti-insulin B cell CD86 expression. n≥ 6 mice per condition, standard deviation is indicated by error bars. *p<0.05 **p<0.01 ***p<0.001 ****P<0.0001 by one-way Anova with Tukey’s multiple comparison.

Addition of mCD22L to the 4-arm PEG modulated this effect, holding B cell proportions at control levels (4-arm PEG-insulin(2)/CD22L(2) 3.40% ±3.05% SD). The insulin-CD22L construct had no effect on B cell proportions (3.11% ± 2.31% SD).

B cells incubated with the 4-arm PEG-insulin(2) construct also had significantly higher proliferation as measured by dye dilution, higher total B cell numbers, higher upregulation of the costimulatory antigen-presenting molecule CD86, and down regulation of the inhibitory molecule CD22 (Figure 5B-E, **Table 1**). These findings suggest that 4-arm PEG-insulin(2) has stimulatory effects on anti-insulin B cells (Figure 5B-E, **Table 1, Figure S2**), likely due to crosslinking and strong signaling via the BCR. The addition of mCD22L to the PEG backbone in 4-arm PEG-insulin(2)/CD22L(2) antagonizes the stimulatory effects of 4-arm PEG-insulin(2), restoring proliferation, total numbers of B cells and levels of CD86 and CD22 expression back to levels comparable to control wells with media only (Figure 5B-E, **Table 1**). Insulin-CD22L did not have any effects on B cell proportion, total number, number of B cells proliferated, or expression of CD86 and CD22 (Figure 5, **Table 1**). WT B cells were not affected by any of the constructs, indicating that the stimulatory effect of the 4-arm PEG-insulin(2) is antigen-specific. (**Figure S3**).

**Table 1:** Average B cell proportion, number and MFI for anti-insulin B cells incubated with insulin constructs.

| Condition | Average Total Live B cells |  | Average B Cell Proportion |  | Average Number of B Cells Proliferated |  | Average CD86 MFI Ratio |  | Average CD22 MFI Ratio |  |
| --- | --- | --- | --- | --- | --- | --- | --- | --- | --- | --- |
|  |  | SD |  | SD |  | SD |  | SD |  | SD |
| Media Only | 2304 | 2130 | 3.44 | 2.53 | 204 | 145 | 0.98 | 0.38 | 1.07 | 0.39 |
| mCD22L-azide | 2397 | 1476 | 3.99 | 2.34 | 144 | 43 | 0.98 | 1.29 | 0.89 | 0.22 |
| 4-arm PEG-Insulin(2) | 13105 | 7393 | 11.99 | 8.24 | 5458 | 3008 | 44.29 | 16.95 | 0.51 | 0.06 |
| 4-arm PEG-Insulin(2)/mCD22L(2) | 2972 | 2819 | 3.4 | 3.05 | 172 | 216 | 6.56 | 6.34 | 0.7 | 0.42 |
| Insulin-mCD22L | 2997 | 2614 | 3.11 | 2.31 | 206 | 243 | 0.94 | 0.54 | 0.72 | 0.16 |

### Effect of Insulin-mCD22L Conjugates on Anti-insulin B cell activation in Response to Anti-CD40

125Tg B6 mouse splenocytes were stimulated with anti-CD40 to simulate T cell help for three days in the presence of insulin conjugates. Insulin-CD22L significantly reduced anti-insulin B cell proliferation by 29% compared to anti-CD40 alone or in the presence of unconjugated mCD22L (51.1% ±11.4% anti-CD40 alone vs 50.5% ± 9.0% anti-CD40 with mCD22L-azide p=0.9926; 51.1% ±11.4% anti-CD40 alone vs 36.3% ±11.8% anti_CD40 with insulin-mCD22L p=0.0313) (Figure 6A, **Table 2**). Once again, the 4-arm PEG-insulin(2) was highly activating, inducing 81.2% more proliferation compared to anti-CD40 alone. The addition of mCD22L to the PEG backbone in 4-arm PEG-insulin(2)/mCD22L(2) again antagonized excess stimulatory effects, returning proliferation to that of B cells stimulated with anti-CD40 alone (51.1%±11.4% anti-CD40 alone vs 46.9%±10.3% anti-CD40 with 4-arm PEG-insulin(2)/mCD22L(2) p=0.6280) (Figure 6B). Anti-CD40 stimulated WT B6 splenocytes were not affected by the insulin constructs (**Figure S4**).

**Figure 6:**
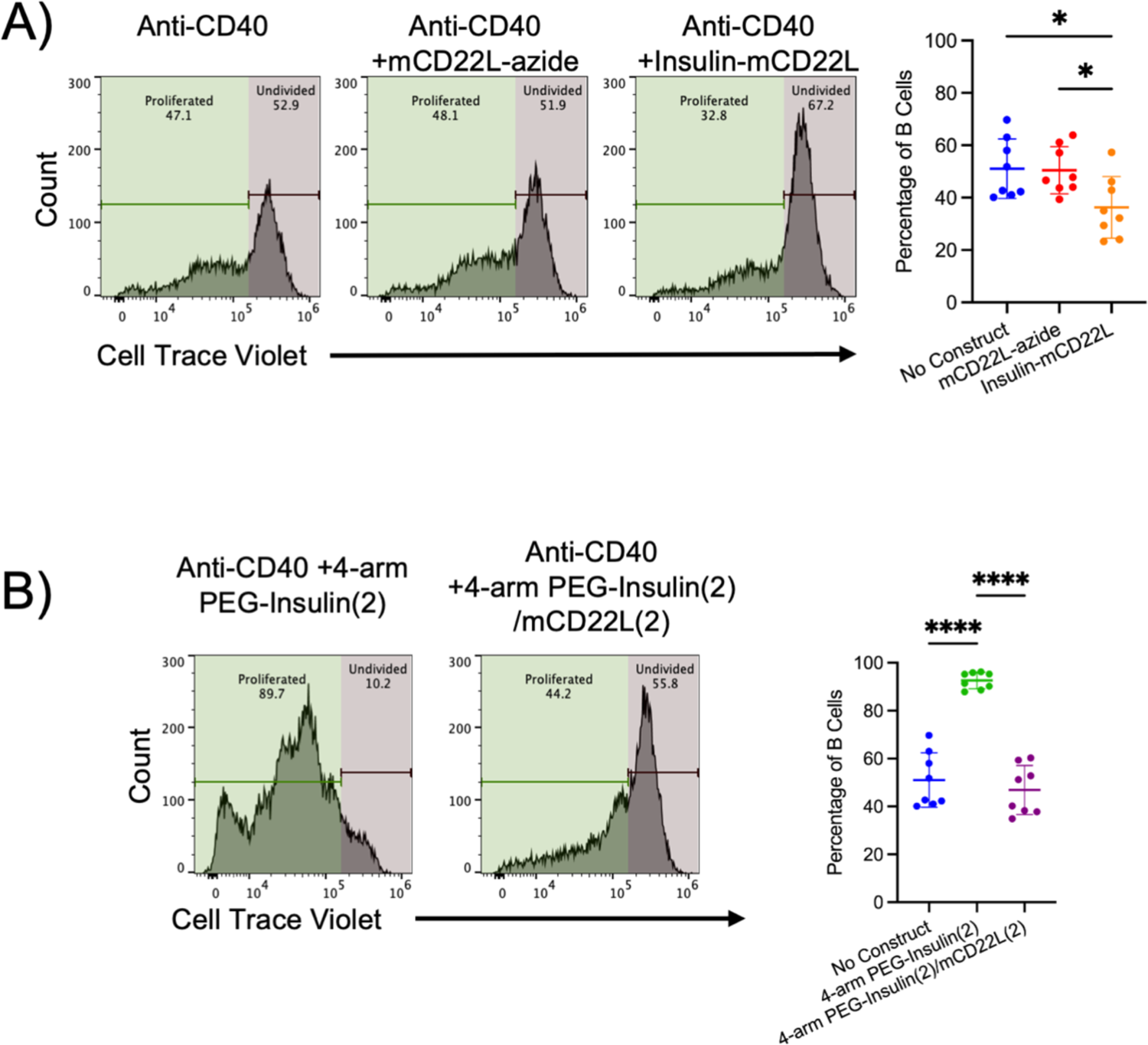
Effects of insulin/mCD22L conjugates on anti-CD40 B cell stimulation in 125Tg B6 mouse splenocytes at 3 days. A) Representative histograms and quantification of B cell proliferation by dye-dilution assay following stimulation with Anti-CD40 and treatment with mCD22L-azide or the insulin-mCD22L direct conjugate (* p<0.05 by one-way Anova with Tukey’s multiple comparison) n:26 mice per condition. B) Representative histograms and quantification of B cell proliferation by dye-dilution assay following stimulation with Anti-CD40 and treatment with 4-arm PEG-insulin(2) or 4-arm PEG-insulin(2)/mCD22L(2) (*p<0.05 **** p<0.0001by one-way Anova with Tukey’s multiple comparison). n=8 mice per condition.

**Table 2:**
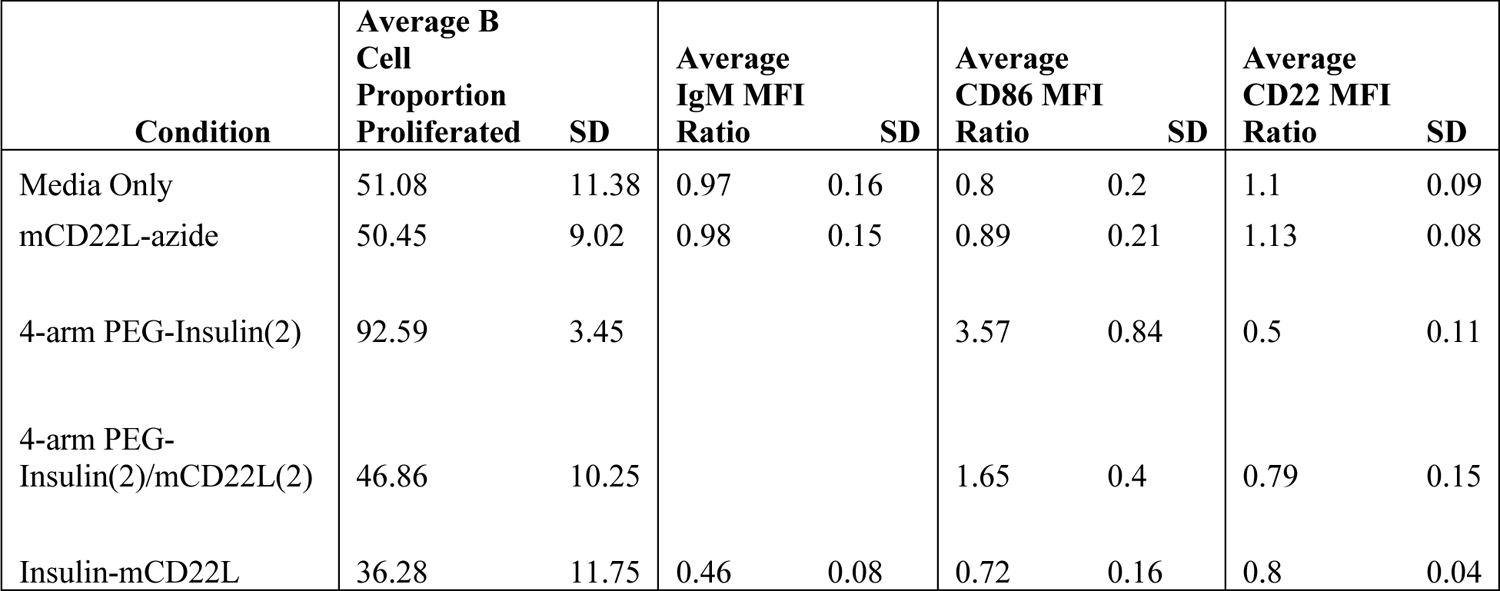
Average Proliferated B cell proportions and MFI for anti-insulin B cells stimulated with anti-CD40 and treated with insulin constructs.

CD86 expression in response to anti-CD40 simulation was indistinguishable from stimulated anti-insulin B cells treated with insulin-mCD22L (Figure 7A, **Table 2**). However, CD86 expression was strongly upregulated in response to anti-CD40 stimulation when treated with 4-arm PEG-insulin(2) and 4-arm PEG-insulin(2)/mCD22L(2) compared with no treatment (Figure 7A). IgM expression was downregulated by 52.4% in B cells stimulated with anti-CD40 when treated with insulin-mCD22L (Figure 7B). Accurate measurement of IgM expression was not feasible with B cells treated with 4-arm PEG-insulin(2) and 4-arm PEG-insulin(2)/ mCD22L(2) potentially due to the treatment occluding the binding of anti-IgM. In WT mice, the insulin conjugates did not significantly affect CD22 or IgM expression, but 4-arm PEG-insulin(2) did increase CD86 expression (**Figure S4**). WT B cell numbers and percent proliferated were not affected (**Figure S4**). CD22 expression was reduced by 26.5% and 28% for 4-arm PEG-insulin(2)/mCD22L(2) and insulin-mCD22L respectively, whereas 4-arm PEG-insulin(2) saw a reduction of 54.7% (Figure 7C).

**Figure 7:**
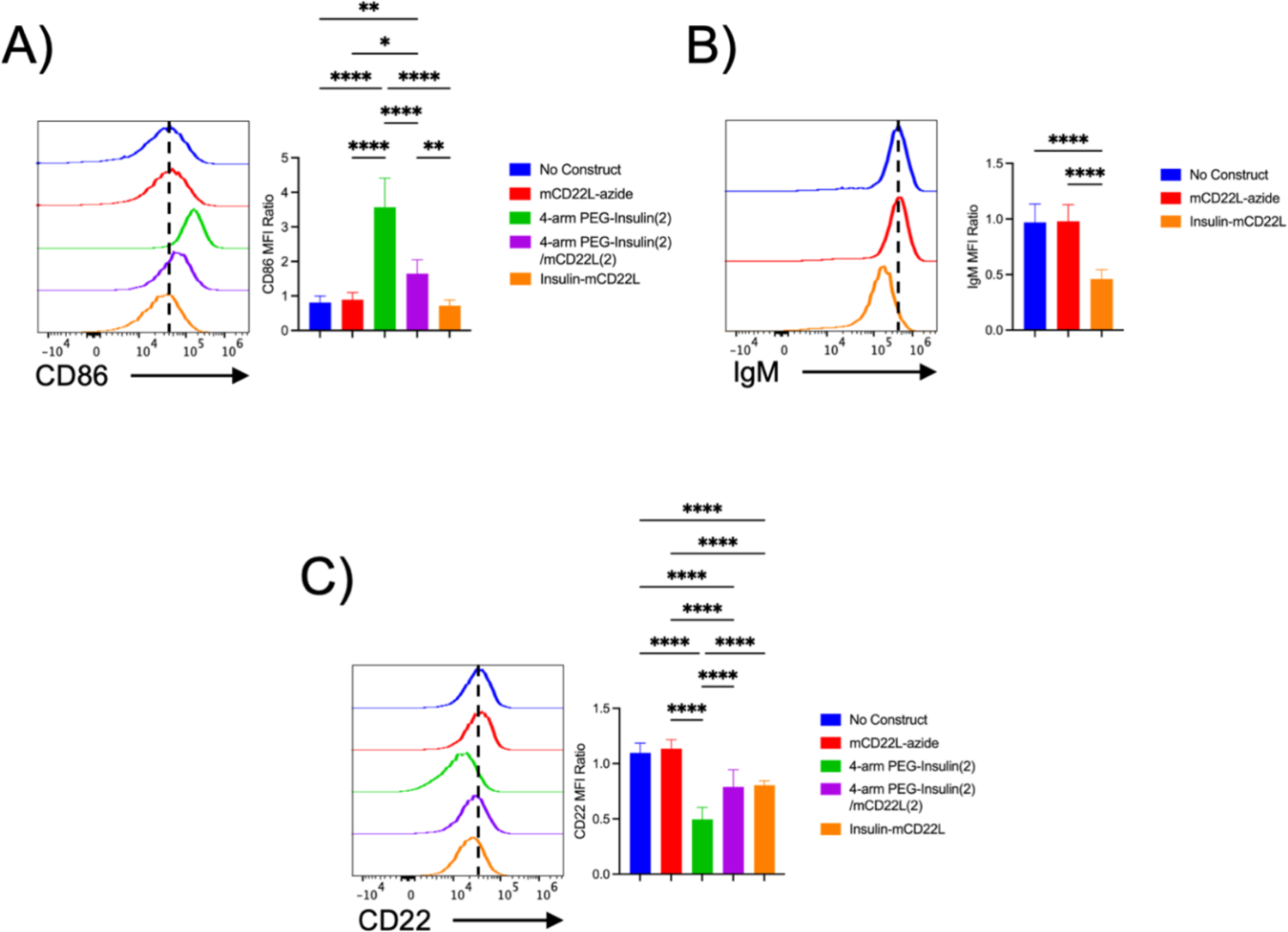
Surface marker expression on anti-insulin B cells after 3 days of anti-CD40 stimulation. A) Representative histograms of CD86 expression levels of anti-insulin B cells and quantification as a ratio over anti-CD40 stimulated WT B cells for treatment with mCD22L-azide, 4-arm PEG-insulin(2), 4-arm PEG-insulin(2)/mCD22L(2) or the insulin-mCD22L direct conjugate following three days of anti-CD40 stimulation. B) Representative histograms of IgM expression levels of anti-insulin B cells and quantification as a ratio over anti-CD40 stimulated WT B cells for treatment with mCD22L-azide or insulin-mCD22L following three days of anti-CD40 stimulation. C) Representative histograms of CD22 expression levels of anti-insulin B cells and quantification as a ratio over anti-CD40 stimulated WT B cells for treatment with unconjugated mCD22L-azide, 4-arm PEG-insulin(2), 4-arm PEG-insulin(2)/mCD22L(2) or the insulin-mCD22L direct conjugate following three days of anti-CD40 stimulation. n:26 mice per condition. Dotted line represents mean MFI of stimulated anti-insulin B cells with no construct. (*p<0.05 ** p<0.01***p<0.001 **** p<0.0001 by one-way Anova with Tukey’s multiple comparison).

## Discussion

### The 4-arm PEG scaffold enables isolation of multivalent insulin-mCD22L conjugates with discrete ligand valencies

Prior studies investigating BCR-CD22 co-ligation have focused on large liposome and polyacrylamide or polynorbornene polymer scaffolds that have antigen and CD22L valences in the range of 100-1000 or more.^3–9^ Additionally, by nature of their synthesis, these scaffolds result in non-discrete ligand valences. Because of the size constraints of liposomes and the heterogeneity of antigen and CD22L valences achieved with linear polymers, a 4-arm PEG-alkyne scaffold was selected to synthesize antigen-CD22L conjugates with discrete ligand valencies to investigate the ability of lower-valency antigen-CD22L conjugates to crosslink BCR and CD22 to prevent B cell activation. Here, the 20 kDa 4-arm PEG-alkyne proved to be a capable scaffold to synthesize and isolate insulin-mCD22L conjugates with discrete ligand valencies as demonstrated by the significant changes in RP-HPLC retention time and mass distributions on MALDI-TOF mass spectra for the intermediate 4-arm PEG-insulin(1-4) products. The increase in product homogeneity is substantial when the analytical data is compared to that of 20 kDa 4-arm PEG-insulin conjugates with different valency prepared by varying the ratio of insulin to 4-arm PEG scaffold.^33^ Although there is 10-20% MW variation in the 20 kDa 4-arm PEG-alkyne, the discrete number of terminal function groups make it a valuable platform to achieve heterofunctional conjugates with discrete ligand valencies.

### The direct insulin-mCD22L conjugate attenuates B cell activation in response to Anti-CD40 stimulation

Treatment of 125Tg mouse splenocytes with insulin-mCD22L significantly reduced B cell proliferation in response to anti-CD40 stimulation. The ability of a monomeric antigen-CD22L construct to limit B cell activation has not been previously demonstrated to our knowledge, and these data support the potential of antigen-CD22L constructs to limit T-dependent B cell activation. Previous studies demonstrated that CD22 knock-out or CD22 mutation to prevent ligand binding result in hyperproliferation following anti-CD40 stimulation in murine B cells, illustrating how CD22 can attenuate CD40 signaling.^36, 37^ The mechanism is not entirely clear, as CD22 does not associate with CD40, and anti-CD40 stimulation alone is incapable of inducing CD22 phosphorylation.^38, 39^ However, anti-CD40 stimulation synergistically increases CD22 ITIM phosphorylation with anti-IgM stimulation via Lyn, further suggesting that CD22 plays a negative feedback role in CD40 activation.^39^ The need for BCR stimulation in tandem with anti-CD40 stimulation to attenuate B cell activation in a CD22-dependent manner is in congruence with our data showing a lack of attenuation when cells were treated with mCD22L-azide. CD40-mediated activating signals are propagated by TRAF1-6 and JAK3 which mediate downstream pathways including the canonical and non-canonical NFkB signaling; p38, Akt and JNK activation; and STAT5 phosphorylation.^40^ These pathways may be attenuated by CD22-activated protein tyrosine phosphatase SHP-1 which can dephosphorylate TRAF3 and associate with TRAF2/6 to regulate downstream signals or by SHP-1 dephosphorylation of downstream effectors such as Syk and BLNK which play a role in both BCR- and CD40-mediated B cell activation.^41–46^ Further study is needed to elucidate the mechanisms by which BCR-CD22 co-ligation attenuates CD40 pro-inflammatory stimulation.

### The addition of 2 mCD22Ls to 4-arm PEG-insulin(2) neutralizes B cell activation

125Tg splenocyte treatment with bivalent 4-arm PEG-insulin(2) increased B cell activation with or without anti-CD40 co-stimulation. The ability of 4-arm PEG-insulin(2) to crosslink BCRs in a fashion similar to anti-IgM Fab2’ is the likely cause of this response, but the large magnitude of B cell activation was unexpected based on prior investigations into the effects of lower-valency insulin-polymer conjugates on insulin-specific B cell activation.^33, 47^ However, the addition of 2 mCD22Ls to 4-arm PEG-insulin(2), to make 4-arm PEG-insulin(2)/mCD22L(2), abrogated the excessive proliferative effects and resulted in CD86 and proliferation levels indistinguishable from those of untreated cells. This also occurred in the presence of anti-CD40 co-simulation. However, unlike the direct insulin-mCD22L conjugate, 4-arm PEG-insulin(2)/mCD22L(2) was not able to reduce B cell activation resulting from anti-CD40 stimulation. Therefore, the inclusion of an equal number of mCD22L with bivalent antigen was only capable of negating the excessive proliferation resulting from BCR cross-linking. In comparison, high-valency antigen and CD22L-bearing liposomes required 10-30-fold more CD22L than antigen to mitigate the inflammatory stimulation of the multi-valent antigen to baseline levels.^5^ BCR and CD40 stimulation is synergistic in B cell activation; therefore, avoiding BCR crosslinking is important to achieve tolerogenic effects with antigen-CD22L constructs in the presence of T cell help.^45, 48, 49^ As demonstrated by prior antigen and CD22L-bearing liposomes, increasing antigen valency may require an exponential increase in CD22L valency to overcome the pro-inflammatory signal resulting from BCR crosslinking.

### Low valency antigen-CD22L conjugates can successfully co-ligate BCR and CD22

High valency antigen-CD22L polymers and liposomes have previously demonstrated the ability of BCR/CD22 co-ligation to attenuate B cell activation, but emphasis has been placed on high CD22L valency being necessary in these constructs to outcompete binding with endogenous cis-ligands of CD22 such as sialic acids present on CD22 molecules.^3–5, 32^ The high-affinity mCD22L used in our insulin-mCD22L conjugates has an approximate K_D_ of 1 μM for mCD22; much higher than the 150 nM concentration of insulin-mCD22L used in the assays.^32^ Additionally, the surface concentration of cis-ligands on the B cells is estimated to be greater than 10 mM although the binding affinity of native CD22Ls is in the 100 μM range.^50^ From these values, it is understood that binding of monovalent CD22L alone to B cells is improbable.

However, conjugation of the CD22L to an antigen possessing sufficient affinity for the BCR can immobilize the antigen-CD22L conjugate on the cell surface through BCR-antigen binding. This mechanism, in which the antigen binds first, results in a high local concentration of the CD22L promoting CD22 binding and BCR/CD22 co-ligation. We postulate the same effect occurs with the 4-arm PEG-insulin(2)/mCD22L(2) although BCR cross-linking also takes place. The exact degree of BCR/CD22 co-ligation achieved by the insulin-mCD22L and the 4-arm PEG-insulin(2)/mCD22L(2) is uncertain, and we agree with the hypothesis that a greater extent of BCR/CD22 co-ligation could be achieved by increasing the CD22L valency of each respective construct, but our data does show that conjugating a single CD22L to a single copy of antigen is capable of BCR/CD22 co-ligation in spite of competition from cis-ligands.

## Conclusion

High valency antigen-CD22L conjugates have previously demonstrated the ability to prevent B cell activation. Here we demonstrated that 1:1 insulin-mCD22L attenuates B cell activation from mimicked T-dependent stimulation. However, a 2:2 insulin-mCD22L conjugate was not able to reduce B cell activation from anti-CD40 stimulation, demonstrating that competing inflammatory and anti-inflammatory signals stemming from BCR crosslinking versus BCR and CD22 co-ligation are considerable. The data suggest that antigen-CD22L constructs possessing a single copy of antigen are well-suited to achieve immunosuppressive effects, making it a promising agent for future preclinical trials.

---

For table of contents only:

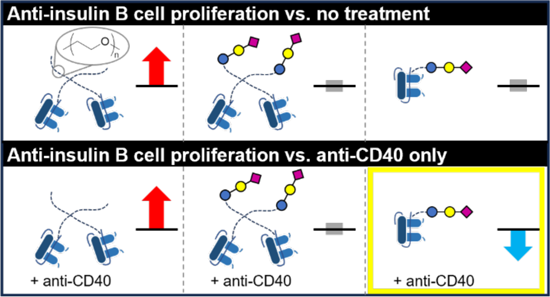

