## Supplemental Information for "CD22L Conjugation to Insulin Attenuates Insulin-Specific B cell Activation"

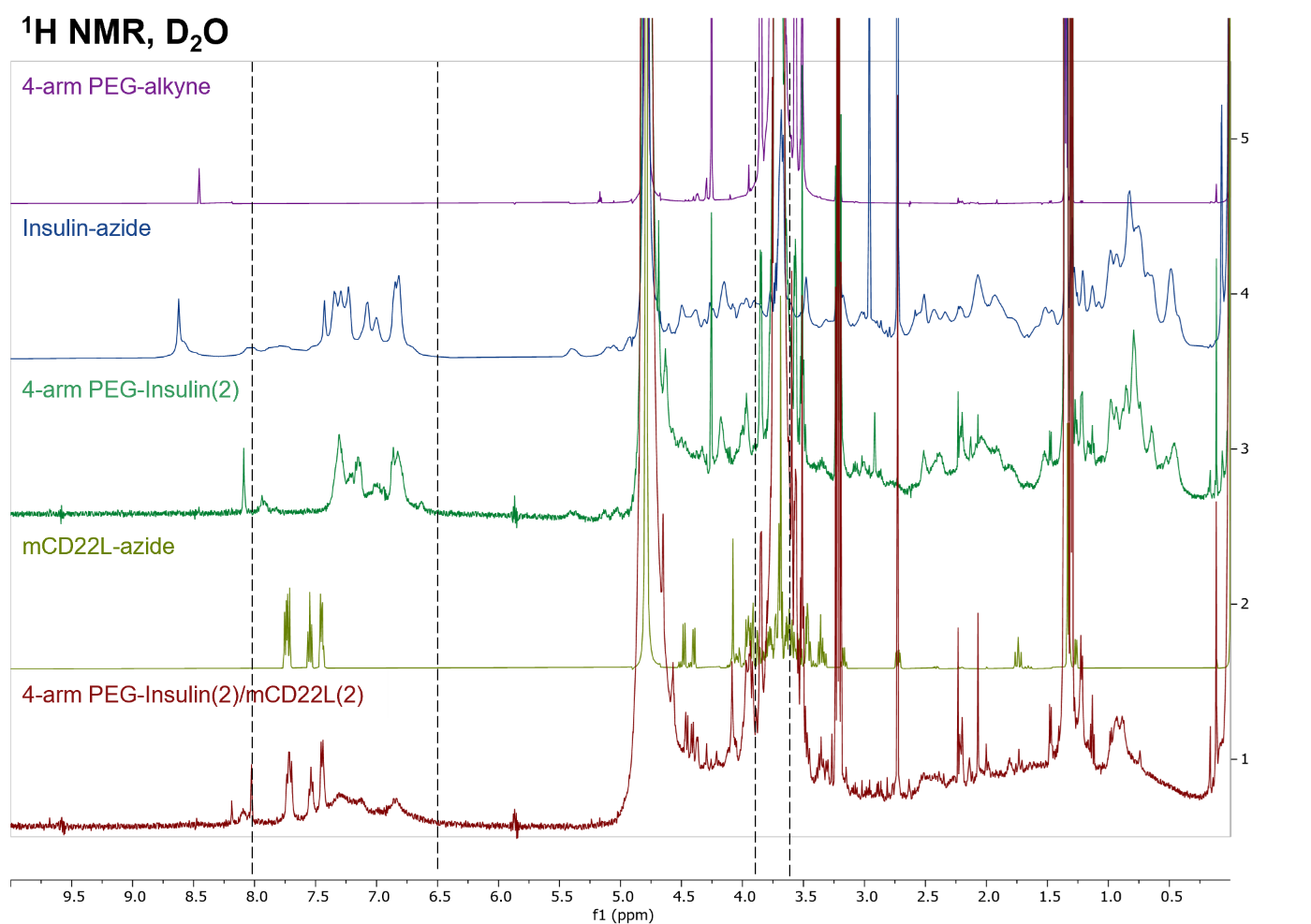

**Figure S1**: Proton NMR spectra of 4-arm PEG-insulin(2)/mCD22L(2) and precursors from 0-10 ppm. Dotted lines indicate the integrated regions for estimating the number of insulin or mCD22L molecules per 4-arm PEG scaffold as detailed in table S1.

**Table S1:** Mass and Proton NMR Integration Data

|  | **MALDI-TOF-MS** | | **Theoretical/Calculated** | | | **Measured** | **Result** |
| --- | --- | --- | --- | --- | --- | --- | --- |
| **Species** | Theoretical MW | Measured MW | Protons (6.5 - 8.0 ppm) | Protons (3.6-3.82 ppm) | (6.5-8.0/3.6-3.82) Ratio | (6.5-8.0/3.6-3.82) Ratio | Measured/ Calculated |
| Insulin-azide | 5,806.7 | 5,807.3 | 35 | 21 | 1.67E+0 | 1.68E+00 | 101% |
| mCD22L-azide | 1,000.0 | 1,000.6 | 9 | 18 | 5.00E-1 | 5.04E-01 | 101% |
| 4-arm PEG (20 kDa) | 20,000 | 20,174.8 | 0 | 1800 | 0.00E+0 | 5.56E-05 | - |
| 4-arm PEG-Insulin(2) | 32,000 | 32,332.3 | 70 | 1828 | 3.83E-2 | 3.16E-02 | 82% |
| 4-arm PEG-insulin(2)/mCD22L(2) | 34,000 | 34,140.8 | 88 | 1864 | 4.72E-2 | 2.34E-02 | 50% |
| ^mCD22L contribution | - | - | 18 | 1864 | 9.66E-3 | 9.01E-03 | 93% |
| ^Insulin contribution | - | - | 70 | 1864 | 3.76E-2 | 1.44E-02 | 38% |

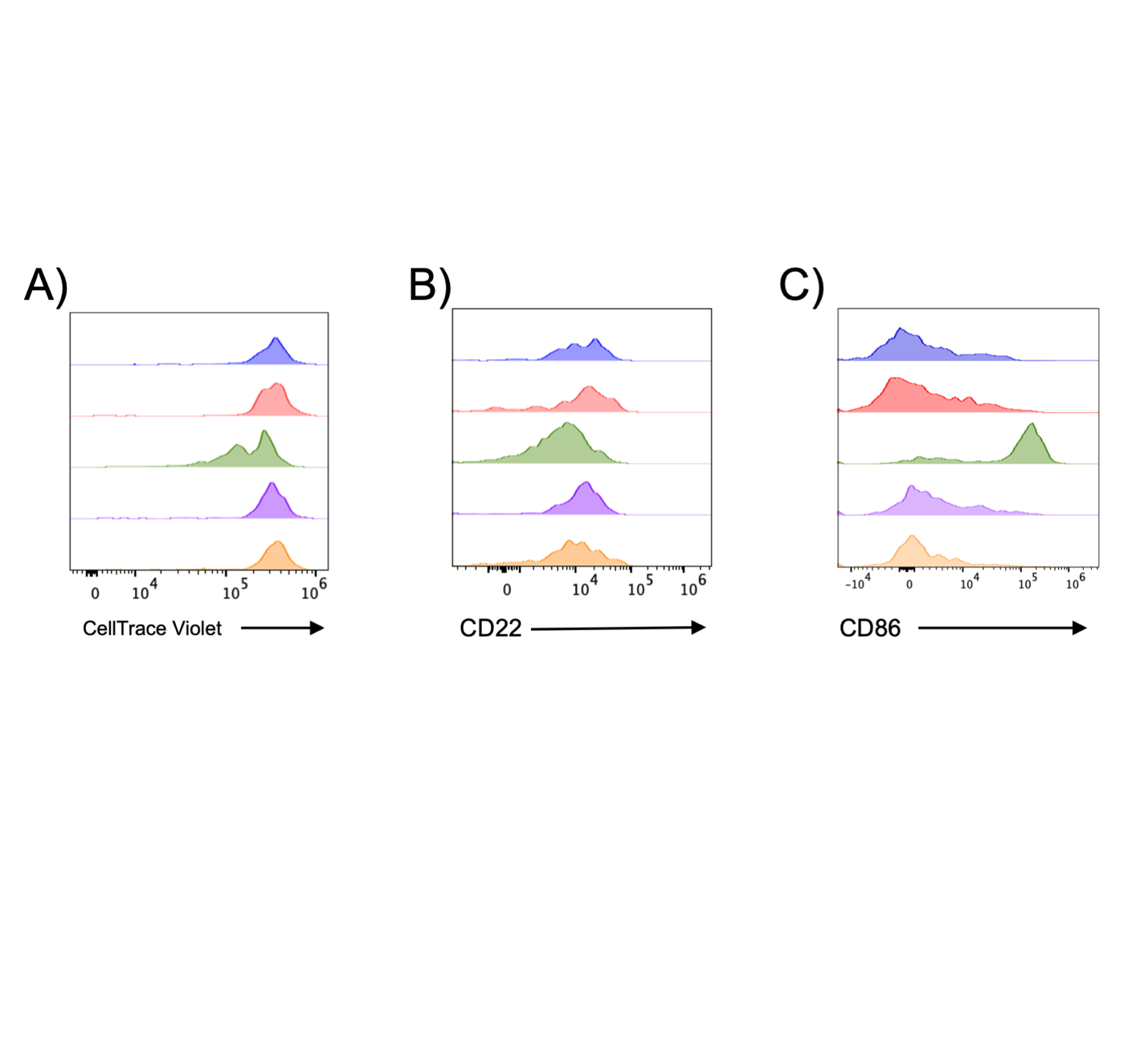

**Figure S2:** Representative histograms of proliferation dilution dye and surface protein expression on 125Tg splenocytes cultured for 72 hours in media (blue), 150nM mCD22L-azide (red), 150nM 4-arm PEG-insulin(2) (green), 150nM 4-arm PEG-insulin(2)/mCD22L(2) (purple), and 150nM insulin-mCD22L (orange). A) Representative of histograms of anti-insulin B cell proliferation by dye dilution assay. B) Representative histogram of surface CD22 expression on anti-insulin B cells. C) Representative histogram of surface CD86 expression on anti-insulin B cells.

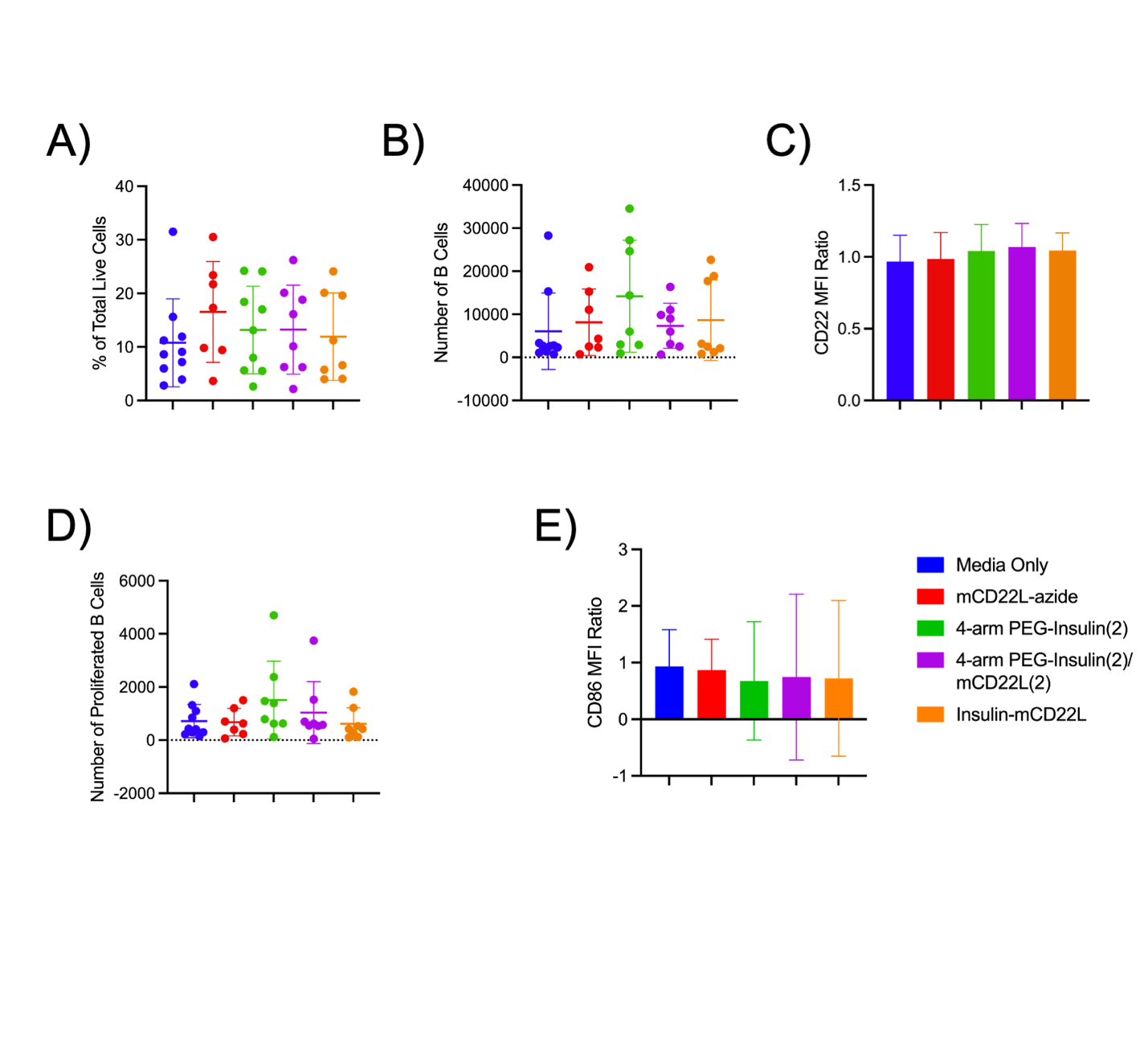

**Figure S3:** WT B cell responses to the incubation of insulin and mCD22L conjugates in B6 mouse splenocytes after 72 hours. A) The percentage of total live cells that are B cells. B) Total live B cell number. C) CD22 expression levels on B cells as a ratio over untreated WT B cell CD22 expression. D) Number of proliferating B cells determined by dye-dilution assay. E) CD86 expression levels on B cells as a ratio over untreated WT B cell CD86 expression. n≥8 mice per condition, standard deviation is indicated by error bars. *p<0.05 **p<0.01 ***p<0.001 ****P<0.0001 by one-way Anova with Tukey’s multiple comparison.
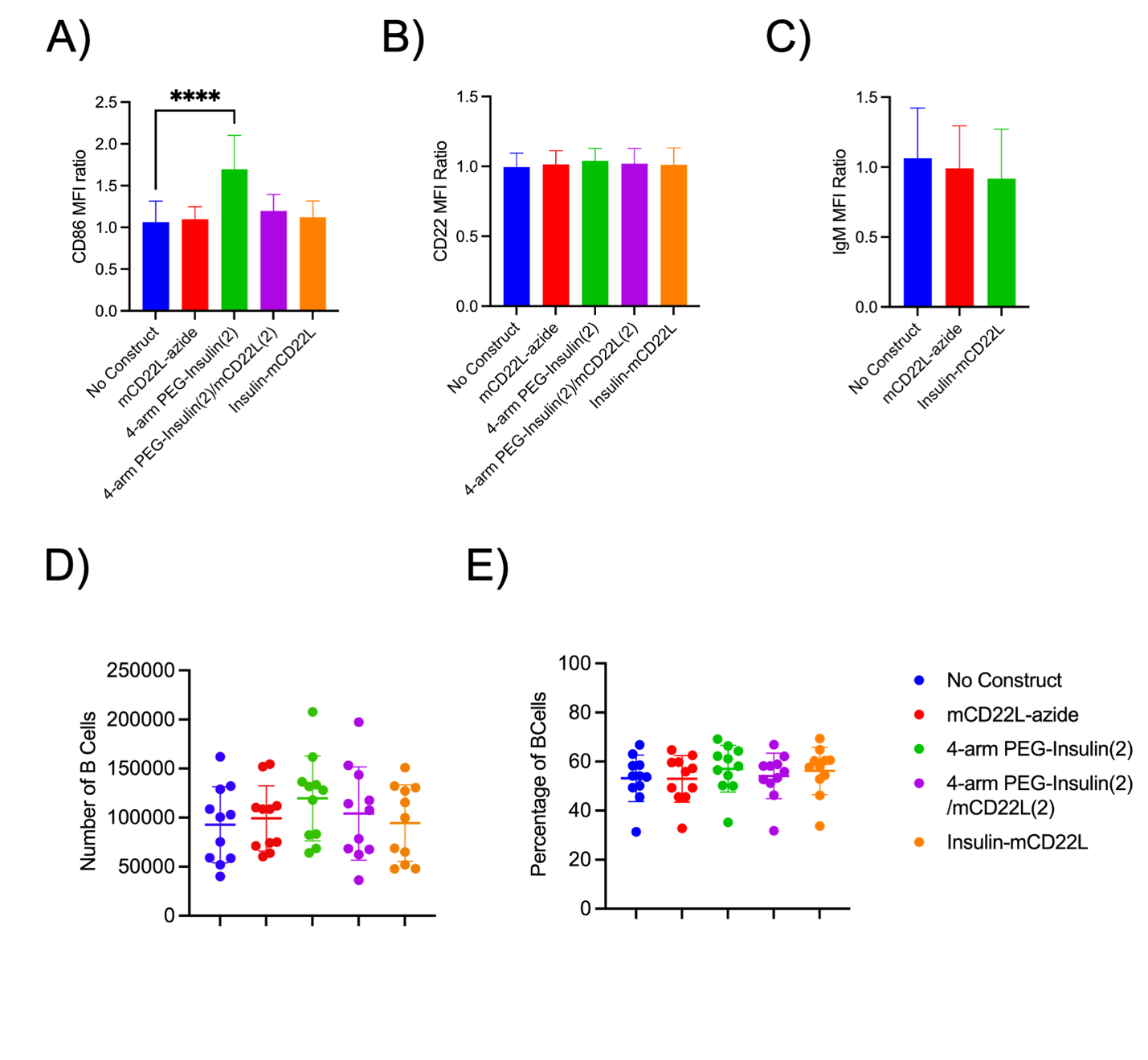

**Figure S4:** Surface marker expression and B cell number of WT splenocytes after 3 days of anti-CD40 stimulation. A) CD86 expression levels as a ratio over anti-CD40 stimulated WT B cells for treatment with mCD22L-azide, 4-arm PEG-insulin(2), 4-arm PEG-insulin(2)/mCD22L(2) or the insulin-mCD22L direct conjugate following three days of anti-CD40 stimulation. B) CD22 expression levels as a ratio over anti-CD40 stimulated WT B cells for treatment with unconjugated mCD22L-azide, 4-arm PEG-insulin(2), 4-arm PEG-insulin(2)/mCD22L(2) or the insulin-mCD22L direct conjugate following three days of anti-CD40 stimulation. C) IgM expression levels as a ratio over anti-CD40 stimulated WT B cells for treatment with unconjugated mCD22L-azide, 4-arm PEG-insulin(2), 4-arm PEG-insulin(2)/mCD22L(2) or insulin-mCD22L following three days of anti-CD40 stimulation. D) Total number of WT B cells following three days of anti-CD40 stimulation. E) B cell proliferation by dye-dilution assay following stimulation with Anti-CD40 and treatment with unconjugated mCD22L-azide 4-arm PEG-insulin(2), 4-arm PEG-insulin(2)/mCD22L(2), or the insulin-mCD22L direct conjugate n≥8 mice per condition (**** p<0.0001 by one-way Anova with Tukey’s multiple comparison).

**Materials and Instrumentation.**

All reactions were carried out in oven- or flame-dried glassware. Commercially obtained chemicals and reagents were used as supplied without further purification unless otherwise stated. Anhydrous solvents were used directly from solvent purification system from innovative technologies. Thin layer chromatography (TLC) was carried out on aluminum backed silica gel TLC plates w/ UV254 (Sorbent Technologies), and revealed by UV irradiation (254 nm) or using CAM, 10% H2SO4 in EtOH or KMnO4 solution followed by developing with gentle heating. Normal phase column chromatography purification of the intermediates and final compounds were performed by a Teledyne ISCO CombiFlash Rf system using appropriate silica gel pre-packed column, and preparative reverse-phase HPLC system Agilent technologies 1210 using preparative column Macherey Nagel HPLC Column Isis 21x250mm, 5 µm. All preparative runs were performed at flow rates of 20.0 ml/min. using a mixture of acetonitrile (HPLC grade, Fisher Chemicals, USA) and Milli-Q-water both acidified with 0.1% TFA. Yields mentioned refer to chromatographically and spectroscopically isolated pure compounds.

NMR spectra were recorded on a 400 MHz spectrometer (Avance III 400), a 500 MHz spectrometer (Avance III 500) with and without cryoprobe from Bruker in MeOD ( 1H, δ = 3.31 ppm, 13C, δ = 49.00 ppm), D2O ( 1H, δ = 4.79 ppm) or CDCl3 ( 1H, δ = 7.26 ppm, 13C, δ = 77.16 ppm) obtained from Cambridge Isotopes Ltd. Chemical shifts (δ) are given in ppm and for the analysis of spectra. The signals were quoted as follows: s = singlet, bs = broad singlet, d = doublet, t = triplet, quin. = quintet, dd = doublet of doublet, ddd = doublet of doublet of doublet, td = triplet of doublet, dt = doublet of triplet and m = multiplet. High-Resolution Mass S9 Spectrometry (HRMS) analysis was performed using an electrospray ion source (ESI) either in positive mode or negative mode and with a time-of-flight (TOF) analyzer on a Waters LCT Premier TM mass spectrometer and are given in m/z. Infrared (IR) spectra were recorded on a Thermo Scientific™ Nicolet™ iS™5 FT-IR Spectrometer; data are reported in frequency of absorption (cm -1); only the most significant adsorption bands are reported; samples are analyzed as thin films deposited on Zn/Se crystal from solvent evaporation.

**Synthesis of mCD22L-azide**

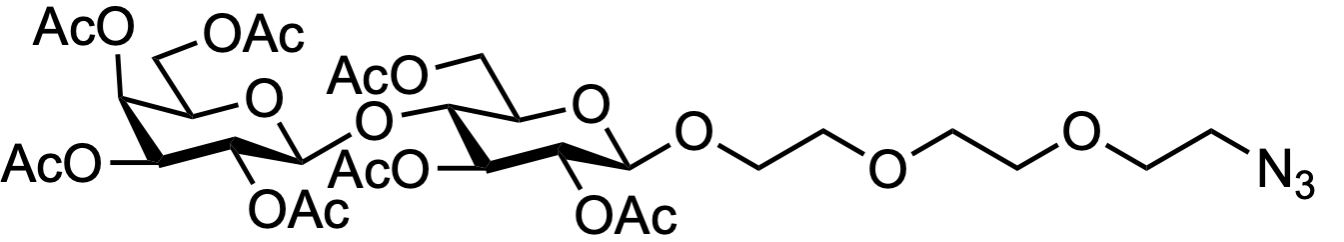
**S1**

A 250 ml round bottom flask was charged with lactose octaacetate (5.0 g,7.37 mmol, 1 equiv.), 2-(2-(2-azidoethoxy)ethoxy)ethan-1-ol (1.94g, 11. 6 mmol, 1.5 equiv.), and CH_2_Cl_2_ (150 ml). The resulting solution was cooled on ice and BF_3_^.^OEt_2_ (1.83 ml, 14.74 mmol, 2 equiv.) was added dropwise. The reaction mixture was allowed to come to room temperature and was stirred overnight. The reaction mixture was poured into saturated aqueous NaHCO_3,_ and the organic layer was separated. The aqueous layer was extracted with CH_2_Cl_2_ (3 x 50 ml), and the combined organic layers dried over Na_2_SO_4_, filtered, and condensed under reduced pressure. The resulting residue was purified by flash chromatography (30-70% EtOAc/Hexanes) to provide **S1** (4.66 g, 5.87 mmol, 80% yield) as a colorless oil. ^1^H NMR (400 MHz, Chloroform-*d*) δ = 5.33 (dd, *J*=3.5, 1.2, 1H), 5.18 (t, *J*=9.3, 1H), 5.09 (dd, *J*=10.4, 7.8, 1H), 4.94 (dd, *J*=10.4, 3.5, 1H), 4.88 (dd, *J*=9.5, 7.9, 1H), 4.56 (d, *J*=7.9, 1H), 4.47 (d, *J*=7.8, 1H), 4.19 – 4.03 (m, 3H), 3.94 – 3.84 (m, 2H), 3.78 (t, *J*=9.4, 1H), 3.74 – 3.56 (m, 13H), 3.38 (dd, *J*=6.9, 3.2, 3H), 2.14 (s, 3H), 2.11 (s, 3H), 2.05 (s, 3H), 2.03 (s, 6H), 2.03 (s, 3H), 1.95 (s, 3H). ^13^C NMR (101 MHz, Chloroform-*d*) δ = 170.2, 170.2, 170.0, 169.9, 169.6, 169.5, 169.0, 100.9, 100.5, 76.1, 72.7, 72.5, 72.4, 71.5, 70.9, 70.5, 70.2, 69.9, 69.0, 68.9, 66.6, 61.9, 61.5, 60.8, 50.5, 20.7(2C), 20.5, 20.5, 20.3. FTIR (thin film): 2873.6, 2102.2, 1747.3, 1368.3, 1220.2, 1047.7, 903.4. HRMS: calc’d [M+Na]^+^ 816.2645, found 816.2661.

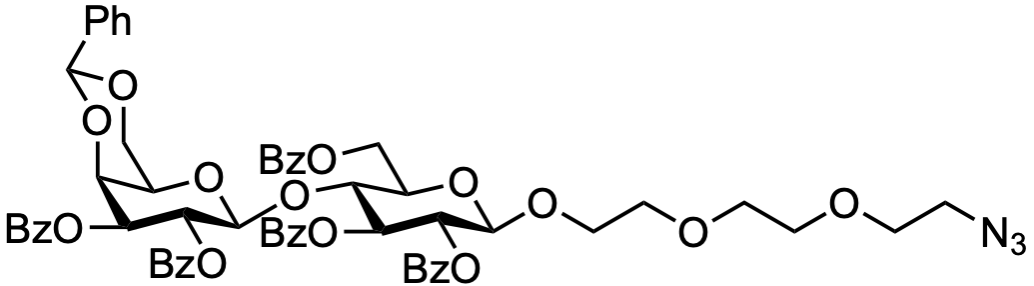
**S2**

An oven-dried 100 ml round bottom flask was charged with **S1** (4.6 g, 6.1 mmol, 1 equiv.), and anhydrous methanol (50 ml). The solution was cooled to 0 °C via ice/water bath, and NaOMe (130 mg, 2.4 mmol, 0.39 equiv.) was added in portions. The resulting solution was allowed to warm to ambient temperature and was stirred until the starting material was consumed (monitored by TLC 20% MeOH/CH_2_Cl_2_). The reaction was quenched with DOWEX H^+^, the suspension was filtered, and the filtrate was condensed under reduced pressure. The resulting residue was dissolved in DMF (50 ml), and benzaldehyde dimethyl acetal (1.4 ml, 9.15 mmol, 1.5 equiv.), and *p*TSA (100 mg, 0.6 mmol, 0.01 equiv.) were added. The reaction was then allowed to stir overnight. The crude reaction mixture was quenched with triethylamine (0.5 ml) and condensed under reduced pressure. The resulting residue was purified by flash chromatography (0-10% MeOH/CH_2_Cl_2_), and fractions containing the desired product were combined, condensed, and directly used in the next step. An oven-dried 50 ml round bottom flask was charged with the crude intermediate and anhydrous pyridine (15 ml). The solution was cooled to 0 °C via ice/water bath, and BzCl (4.2 ml, 36 mmol, 6 equiv.) was added dropwise. The resulting solution was allowed to warm to ambient temperature and was stirred overnight. The reaction was then quenched with MeOH (4 ml) and condensed under reduced pressure. The resulting residue was purified by flash chromatography (0-15% acetone/toluene) to provide **S2** (3.0 g, 2.7 mmol, 44% yield) as a colorless solid. ^1^H NMR (400 MHz, Chloroform-*d*) δ = 8.15 – 8.07 (m, 0.4H), 8.05 – 7.98 (m, 2H), 7.98 – 7.93 (m, 2H), 7.93 – 7.86 (m, 6H), 7.64 – 7.53 (m, 1H), 7.54 – 7.41 (m, 6H), 7.41 – 7.27 (m, 12H), 7.17 (t, *J*=7.6, 2H), 5.86 (t, *J*=9.1, 1H), 5.81 (dd, *J*=10.4, 7.9, 1H), 5.34 (dd, *J*=9.5, 7.8, 1H), 5.30 (s, 1H), 5.18 (dd, *J*=10.4, 3.5, 1H), 4.87 (d, *J*=7.9, 1H), 4.81 (d, *J*=7.8, 1H), 4.64 (dd, *J*=12.0, 2.1, 1H), 4.39 (dd, *J*=12.1, 4.2, 1H), 4.32 (d, *J*=3.7, 1H), 4.24 (t, *J*=9.3, 1H), 3.90 – 3.83 (m, 2H), 3.79 (dd, *J*=12.4, 1.5, 1H), 3.69 (ddd, *J*=11.2, 7.1, 3.6, 1H), 3.59 (dd, *J*=12.6, 2.0, 1H), 3.56 – 3.46 (m, 4H), 3.44 – 3.38 (m, 2H), 3.34 (dd, *J*=5.6, 3.4, 2H), 3.28 (t, *J*=5.1, 2H), 2.98 (s, 1H). ^13^C NMR (101 MHz, CDCl_3_) δ 166.2, 165.7, 165.4, 165.1, 164.9, 137.5, 133.6, 133.4, 133.2, 133.0, 130.2, 129.9 x 2, 129.8 x 2, 129.7, 129.6 x 2, 129.1, 129.0, 128.9, 128.8, 128.5, 128.5, 128.4, 128.4, 128.3, 128.3, 128.0, 126.4, 101.5, 100.9, 100.7, 77.4, 77.1, 76.8 x 2, 74.1, 73.1, 72.9, 72.6, 72.4, 70.7, 70.5, 70.4, 69.9, 69.5, 69.3, 68.0, 66.5, 62.4, 50.6. FTIR (thin film): 3055.4, 2865.0, 2105.3, 1727.7, 1451.4, 1315.0, 1270.8, 1177.2, 1096.0, 1069.9, 1026.9, 710.0. HRMS: calc’d [M+Na]^+^1130.3529, found 1130.3552.

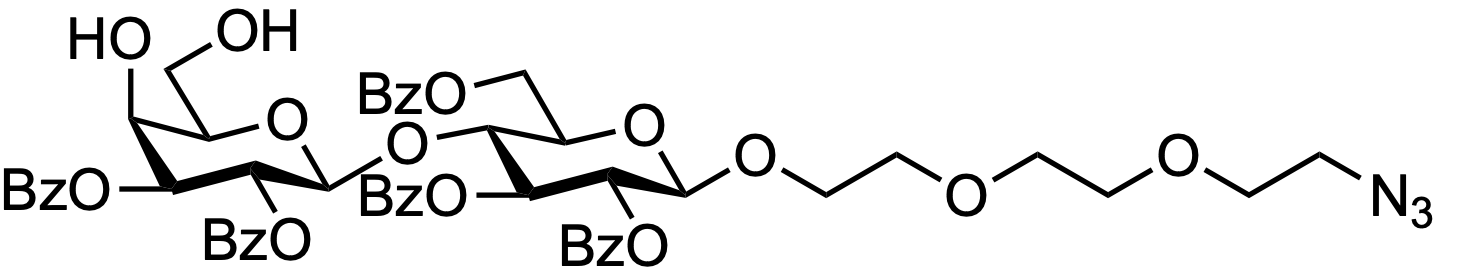
**S3**

A 10 ml round bottom flask was charged with **S2** (100 mg, 0.09 mmol, 1 equiv.), CH_2_Cl_2_ (1 ml) and a few drops of water. The solution was cooled to 0 °C and trifluoracetic acid (0.1 ml) was added. The solution was allowed to warm to ambient temperature and stir overnight. The reaction was poured into sat. NaHCO_3_ and the organic layer was separated. The aqueous layer was further extracted with CH_2_Cl_2_ (3 x 2 ml), dried over Na_2_SO_4_, filtered, and concentrated. The resulting solid was then purified by flash chromatography (0-20% acetone/toluene) to provide **S3** (65 mg, 0.64 mmol, 70% yield) as a colorless solid. ^1^H NMR (400 MHz, Chloroform-*d*) δ = 8.06 – 7.98 (m, 2H), 7.98 – 7.92 (m, 4H), 7.91 – 7.85 (m, 4H), 7.61 – 7.54 (m, 1H), 7.54 – 7.48 (m, 2H), 7.48 – 7.34 (m, 7H), 7.34 – 7.27 (m, 3H), 7.24 – 7.17 (m, 2H), 5.75 (d, *J*=10.8, 1H), 5.73 (d, J=9.6, 1H), 5.40 (dd, *J*=9.5, 7.8, 1H), 5.09 (dd, *J*=10.4, 3.1, 1H), 4.80 (dd, *J*=9.6, 7.8, 2H), 4.59 (dd, *J*=12.1, 2.0, 1H), 4.41 (dd, *J*=12.1, 5.0, 1H), 4.23 – 4.13 (m, 2H), 3.93 – 3.80 (m, 2H), 3.69 (ddd, *J*=11.2, 7.0, 3.7, 1H), 3.58 – 3.50 (m, 2H), 3.50 – 3.45 (m, 2H), 3.44 – 3.30 (m, 6H), 3.30 – 3.30 (m, 3H). ^13^C NMR (101 MHz, CDCl_3_) δ 165.9, 165.8, 165.6, 165.3, 165.2, 133.5, 133.4, 133.3, 133.2, 129.9, 129.7, 129.7 x 2, 129.5, 129.1, 128.9, 128.7, 128.5, 128.4, 101.4, 100.9, 77.4, 77.1, 76.8, 76.6, 74.4, 74.3, 73.7, 73.0, 72.0, 70.7, 70.5, 70.4, 69.9, 69.3, 67.9, 62.7, 62.1, 50.6. FTIR (thin film): 2927.1, 2865.3, 1726.2, 1273.9, 1069.7, 711.5. HRMS: calc’d [M+Na]^+^1042.3219, found 1042.3223.

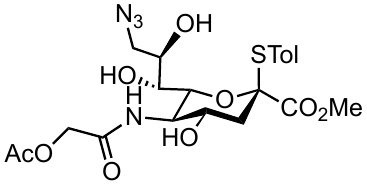
 **S4**
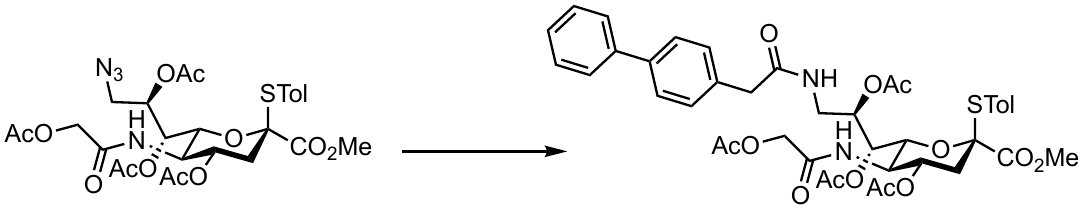
**S5**

A 250 ml 2 neck flask was charged with thioglycoside **S4**^1^ (1.3 g, 3.34 mmol, 1 equiv.) and pyridine/H_2_O/triethylamine (5:2:1, 190 ml). The resulting solution was cooled on ice, and H_2_S bubbled into the reaction flask for ~2 hours. The reaction mixture was stirred at room temperature for ~4 hours, and starting material was consumed. The reaction was then condensed. The crude reaction mixture was dissolved in DMF (50 ml), and diisopropylethylamine (2 ml) and 2,5-dioxopyrrolidin-1-yl 2-([1,1'-biphenyl]-4-yl)acetate were added. The reaction was then allowed to stir overnight. The reaction was then condensed, and the resulting oil was then purified by flash chromatography (5% MeOH/CH_2_Cl_2_) fractions containing desired material were collected and directly used for the next step. A 25 ml round bottom flask containing the crude material was dissolved in pyridine (4 ml). The resulting solution was cooled on ice, and Ac_2_O (960 µl, 10.14 mmol, 30 eq.) was added dropwise. The reaction mixture was stirred overnight. The reaction was then quenched with MeOH (2 ml) and then condensed under reduced pressure. The resulting oil was then purified by flash chromatography (70-90% EtOAc/Hexanes) to provide **S5** (260 mg, 0.38 mmol, 11% yield) as a yellow solid. ^1^H NMR (500 MHz, CDCl_3_) δ 7.55 (td, *J* = 7.6, 1.6 Hz, 4H), 7.42 (t, *J* = 7.7 Hz, 2H), 7.38 – 7.30 (m, 5H), 7.15 (d, *J* = 7.9 Hz, 2H), 6.27 (t, *J* = 6.2 Hz, 1H), 6.07 (d, *J* = 10.2 Hz, 1H), 5.43 (td, *J* = 11.2, 4.7 Hz, 1H), 5.26 (dd, *J* = 3.8, 2.6 Hz, 1H), 4.89 (q, *J* = 4.5 Hz, 1H), 4.68 (dd, *J* = 10.5, 2.6 Hz, 1H), 4.57 (d, *J* = 15.3 Hz, 1H), 4.27 (d, *J* = 15.4 Hz, 1H), 4.16 (t, *J* = 10.4 Hz, 1H), 4.12 (q, *J* = 7.2 Hz, 1H), 3.76 (ddd, *J* = 14.7, 6.7, 4.7 Hz, 1H), 3.67 (s, 3H), 3.44 (d, *J* = 2.2 Hz, 2H), 3.17 (dt, *J* = 14.7, 5.3 Hz, 1H), 2.60 (dd, *J* = 13.8, 4.8 Hz, 1H), 2.34 (s, 3H), 2.20 (s, 3H), 2.07 (s, 3H), 2.05 – 2.01 (m, 1H), 2.00 (s, 3H), 1.98 (s, 3H), 1.27 – 1.23 (m, 1H). ^13^C NMR (126 MHz, CDCl_3_) δ 171.16, 170.96, 170.88, 170.81, 169.85, 168.42, 167.82, 140.79, 140.71, 140.07, 136.44, 134.53, 130.20, 129.68, 128.91, 127.49, 127.44, 127.11, 125.28, 88.91, 72.92, 72.86, 69.36, 68.33, 62.80, 52.91, 49.61, 43.23, 38.76, 37.26, 21.46, 21.12, 20.97, 20.86, 20.83. FTIR (thin film): 1737.8, 1655.4, 1531.6, 1369.7, 1219.7, 1157.9, 1098.6, 1032.3, 729.8, 503.4. HRMS: calc’d [M+Na]^+^ 703.2296, found 703.2302.

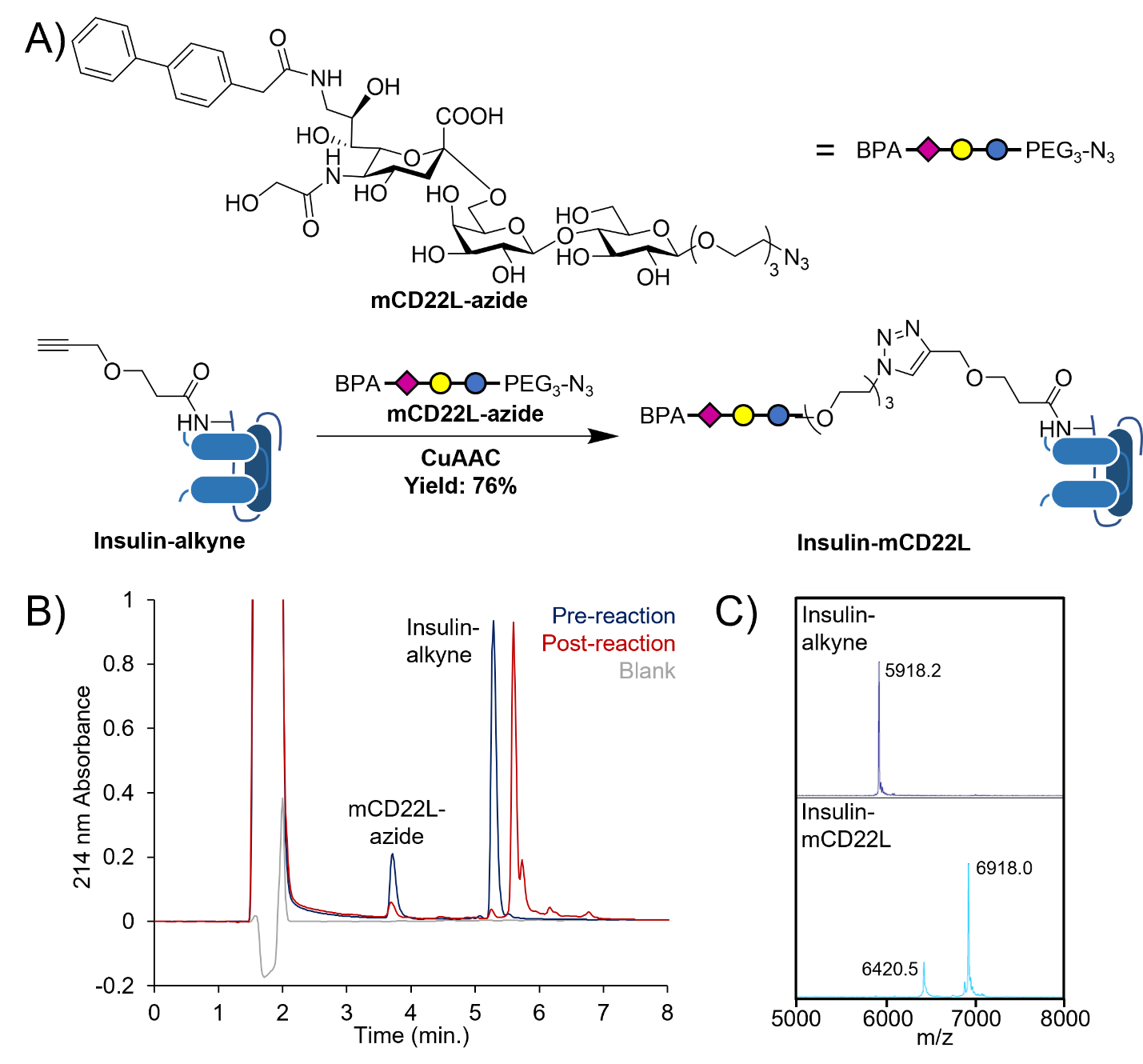

A 100 ml 3 neck flask was charged with thioglycoside **S5** (223 mg, 0.27 mmol, 1.2 equiv.), glycosyl acceptor **S3** (234 mg, 0.23 mmol, 1 equiv.), 4Å molecular sieves (100 mg) and acetonitrile/CH_2_Cl_2_ (5 ml, 4:1). The resulting suspension was stirred overnight, then cooled to -50 °C, and NIS (130 mg, 0.575 mmol, 2.5 equiv.) and TfOH (8 µl, 0.92 mmol, 0.4 equiv.). The reaction was then warmed to -40 °C, stirred for ~20 hours, quenched with triethylamine filtered, and condensed under reduced pressure. The crude reaction mixture was purified by flash chromatography (3% MeOH/CH_2_Cl_2_), and all fractions containing desired trisaccharide were combined and used in the next step as a mixture of ⍺- and β-anomers. A 10 ml round bottom flask containing a mixture of ⍺- and β-anomers was dissolved in MeOH (2 ml). The resulting solution was cooled on ice, and NaOMe (10 mg) was added. The solution was then warmed to room temperature and stirred for ~6 hours. The solution was then neutralized with DOWEX H^+^ to pH 7 filtered and condensed under reduced pressure. The crude reaction mixture was then dissolved in 0.05 M NaOH and stirred overnight. The reaction mixture was then quenched with DOWEX H^+^ filtered and condensed under reduced pressure. The crude reaction mixture was purified by prep-HPLC to provide **mCD22L-azide** (394 mg, 0.34 mmol, 30%) as a colorless solid. ^1^H NMR (400 MHz, D_2_O) δ 7.62 (td, *J* = 8.7, 4.2 Hz, 5H), 7.44 (t, *J* = 7.6 Hz, 2H), 7.35 (t, *J* = 7.4 Hz, 4H), 4.37 (d, *J* = 8.1 Hz, 1H), 4.30 (d, *J* = 7.8 Hz, 1H), 3.99 (s, 2H), 3.97 – 3.78 (m, 6H), 3.79 – 3.64 (m, 4H), 3.64 – 3.41 (m, 14H), 3.40 – 3.32 (m, 3H), 3.30 – 3.20 (m, 2H), 2.64 (dd, *J* = 12.5, 4.7 Hz, 1H), 1.65 (t, *J* = 12.2 Hz, 1H). ^13^C NMR (101 MHz, D_2_O) δ 175.67, 174.48, 173.46, 139.94, 139.10, 134.53, 129.77, 129.11, 127.67, 127.12, 126.75, 103.20, 101.99, 100.46, 79.52, 74.63, 74.58, 73.59, 72.75, 72.43, 72.19, 70.83, 70.09, 69.63, 69.53, 69.44, 69.15, 68.60, 68.52, 68.08, 63.47, 61.06, 60.32, 51.56, 50.10, 42.58, 42.06, 40.20. FTIR (thin film) 3303.0, 2106.7, 1613.9, 1540.1, 1299.6, 1032.0, 821.1, 756.9, 697.8, 556.5. HRMS: calc’d [M+Na]^+^ 1022.3700, found 1022.3696.

**NMR Spectra for mCD22L-azide synthesis**

**S1 ^1^H NMR**

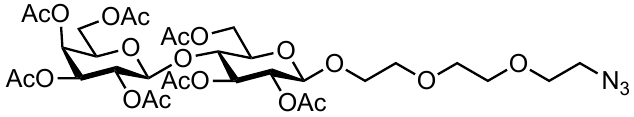

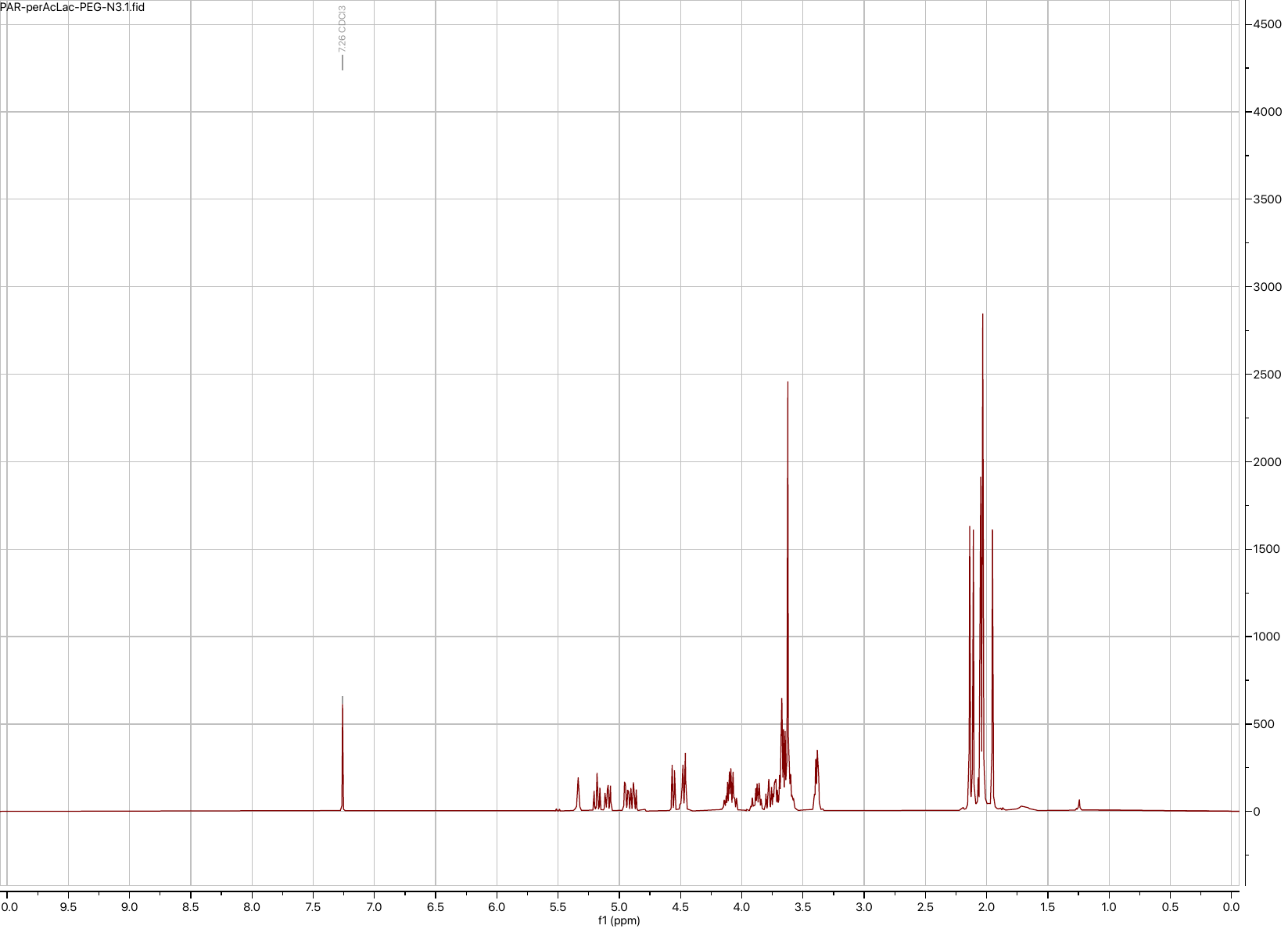

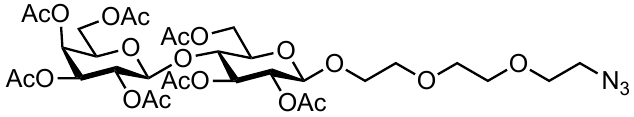

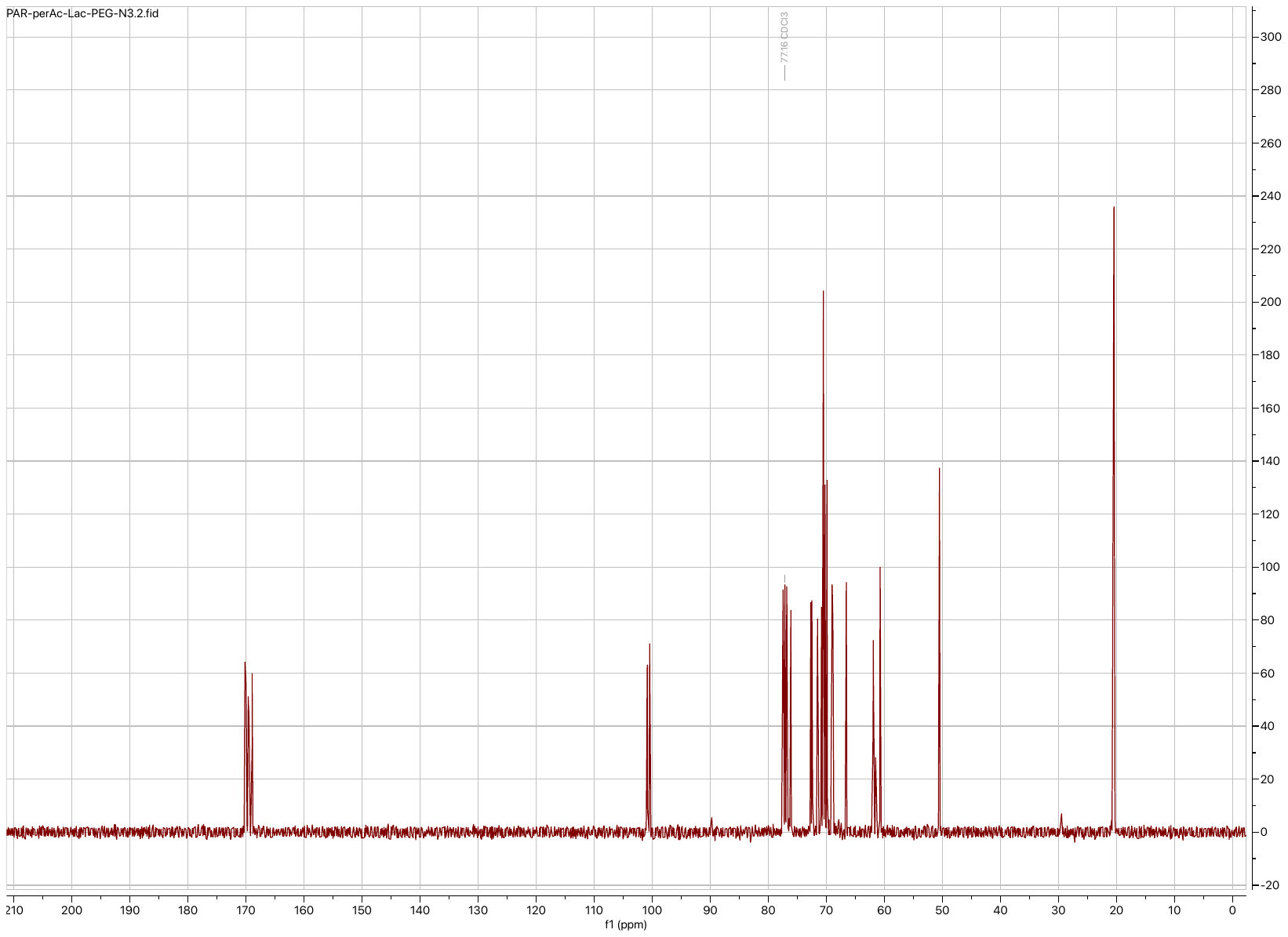

**S1 ^13^C NMR**

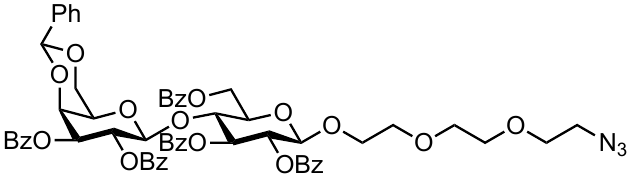

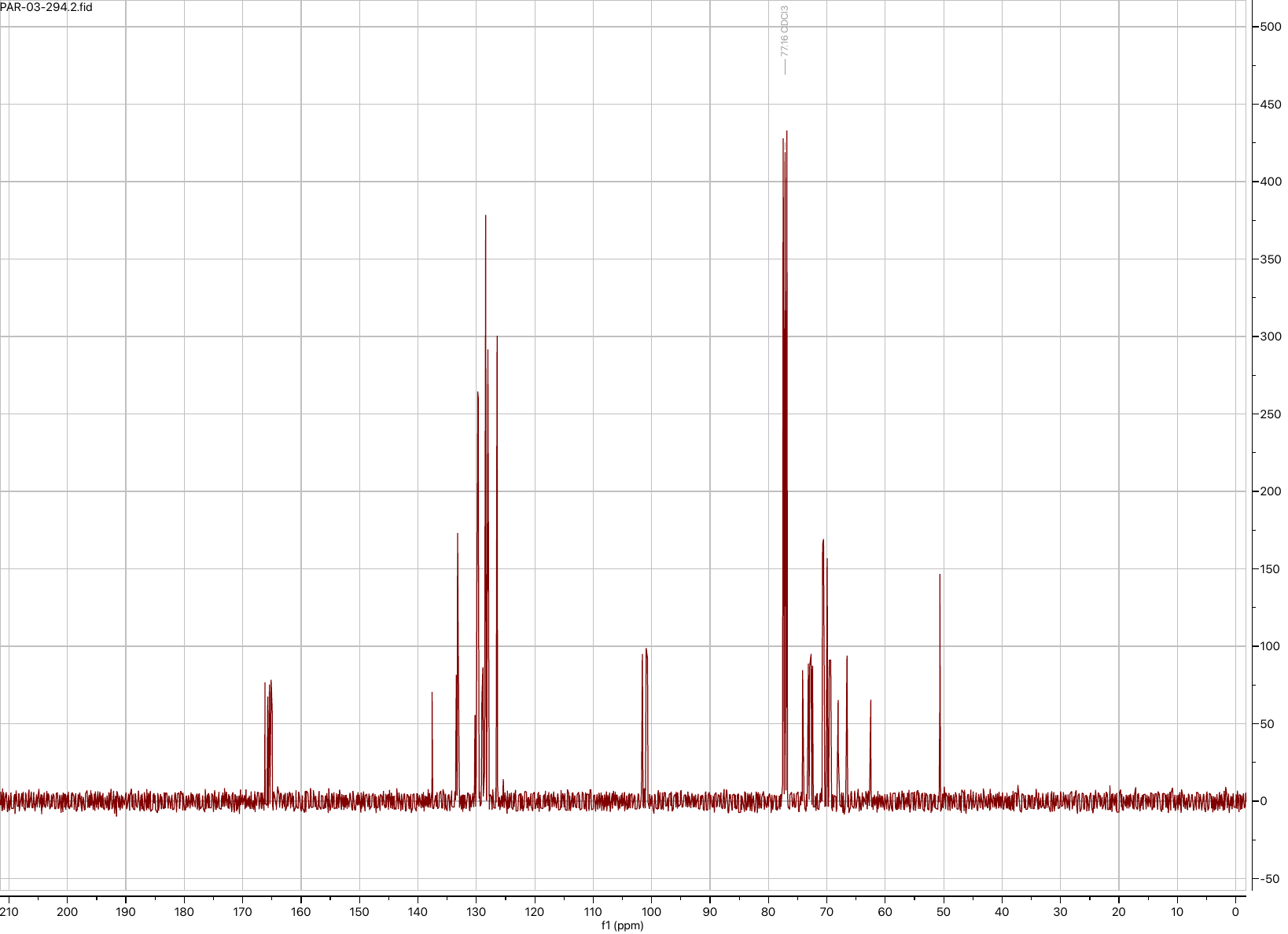

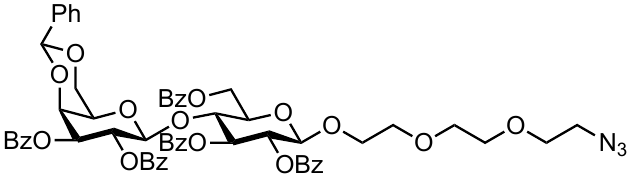

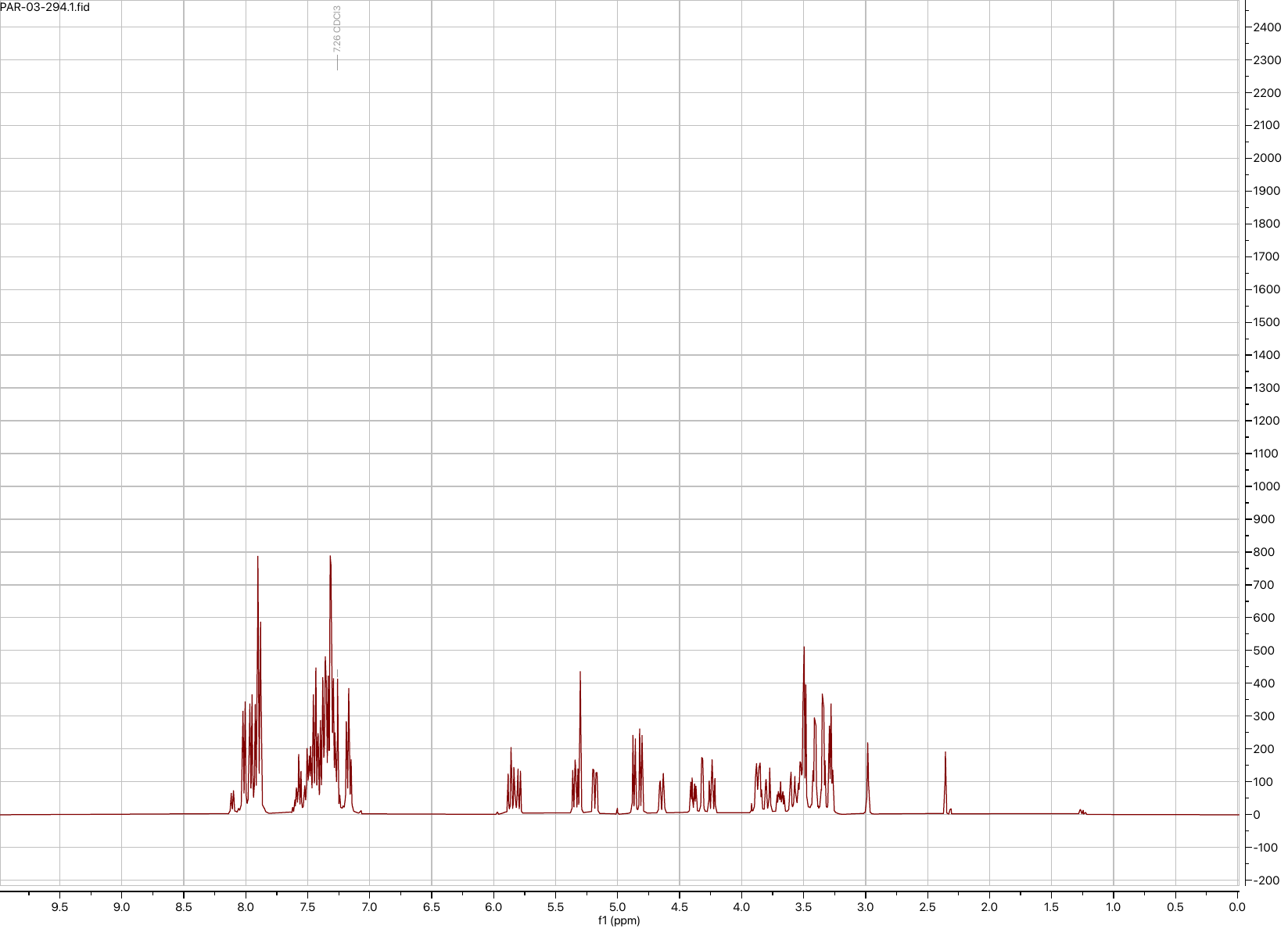

**S2 ^1^H NMR**

**S2 ^13^C NMR**

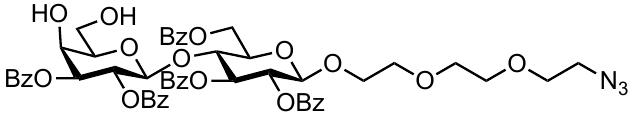

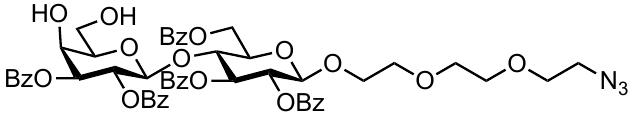

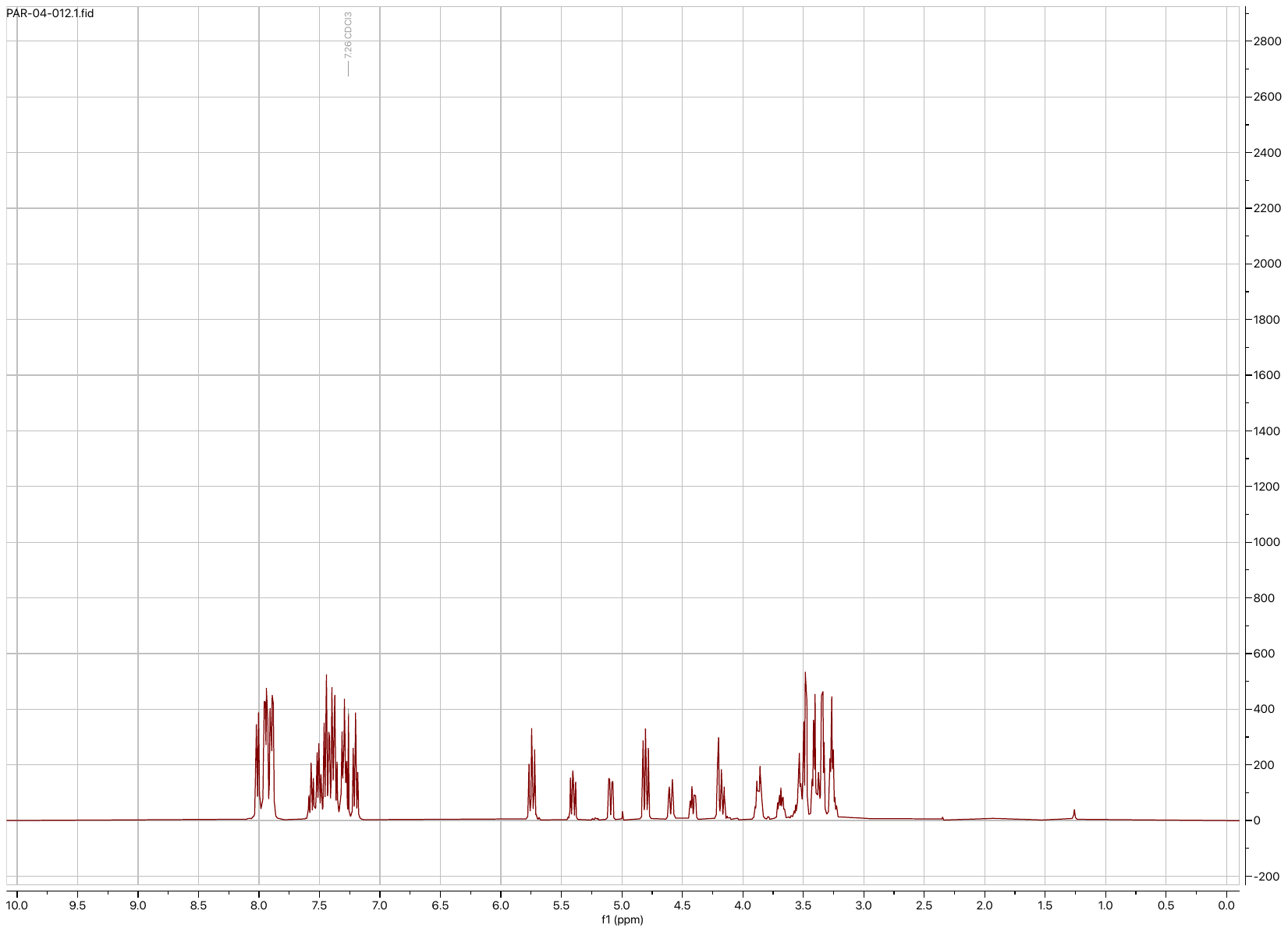

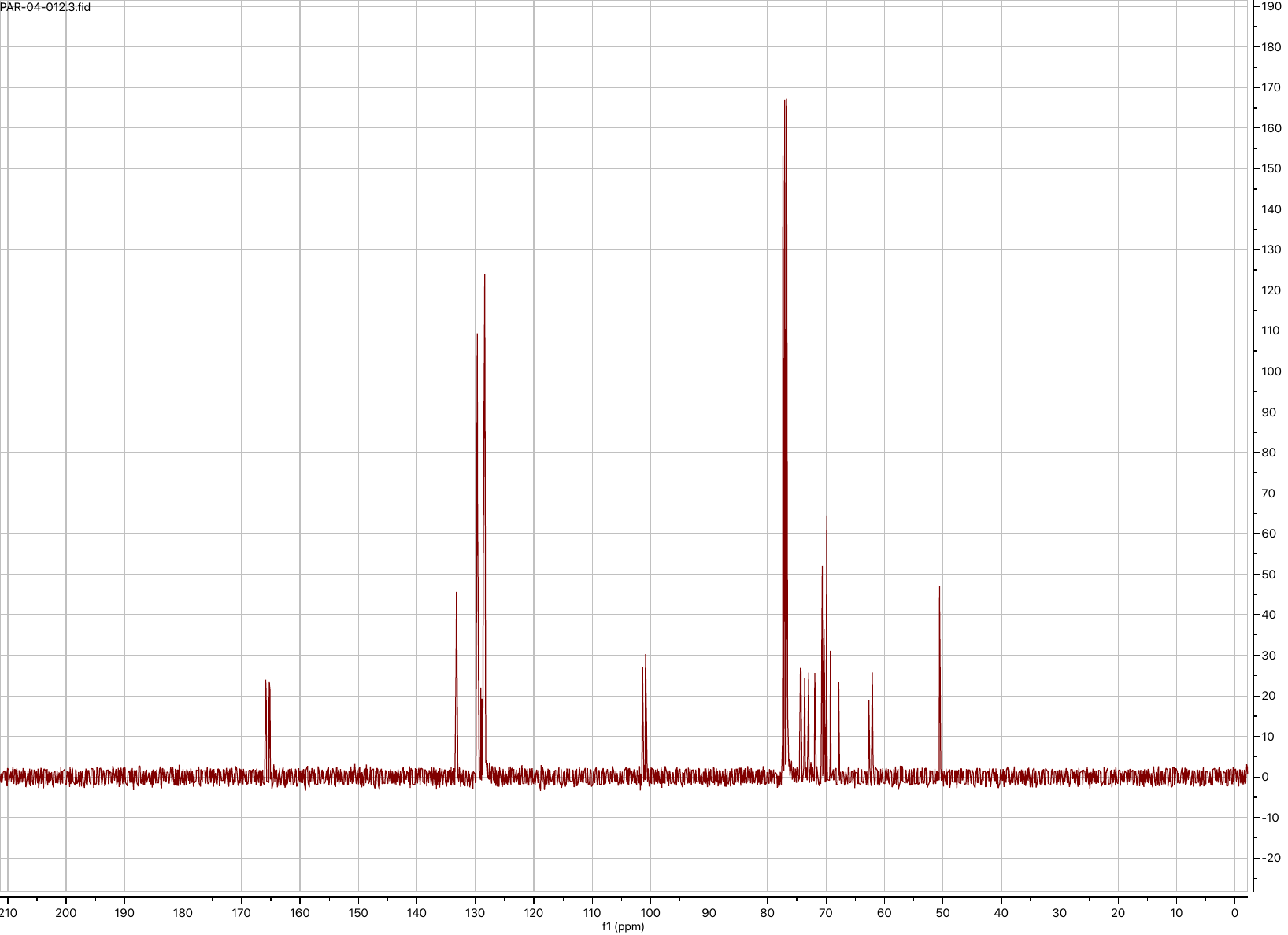

**S3 ^13^C NMR**

**S3 ^1^H NMR**

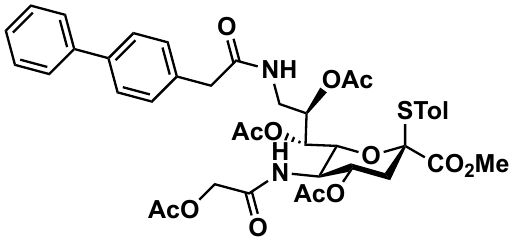

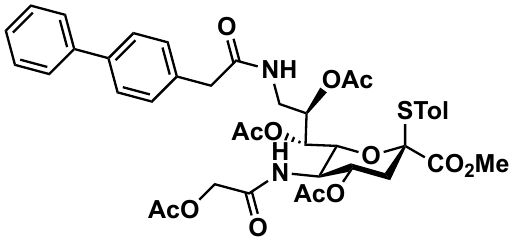

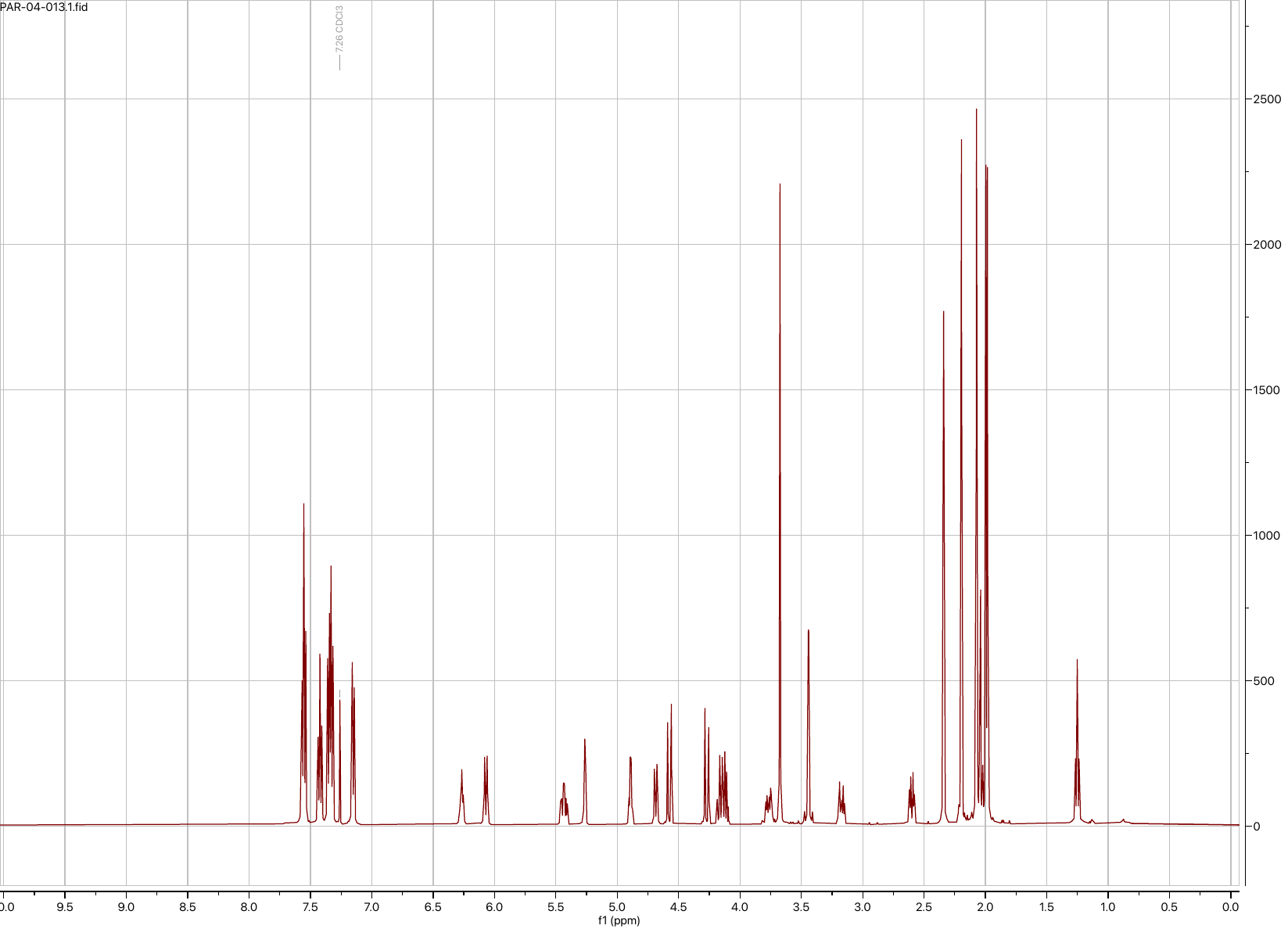

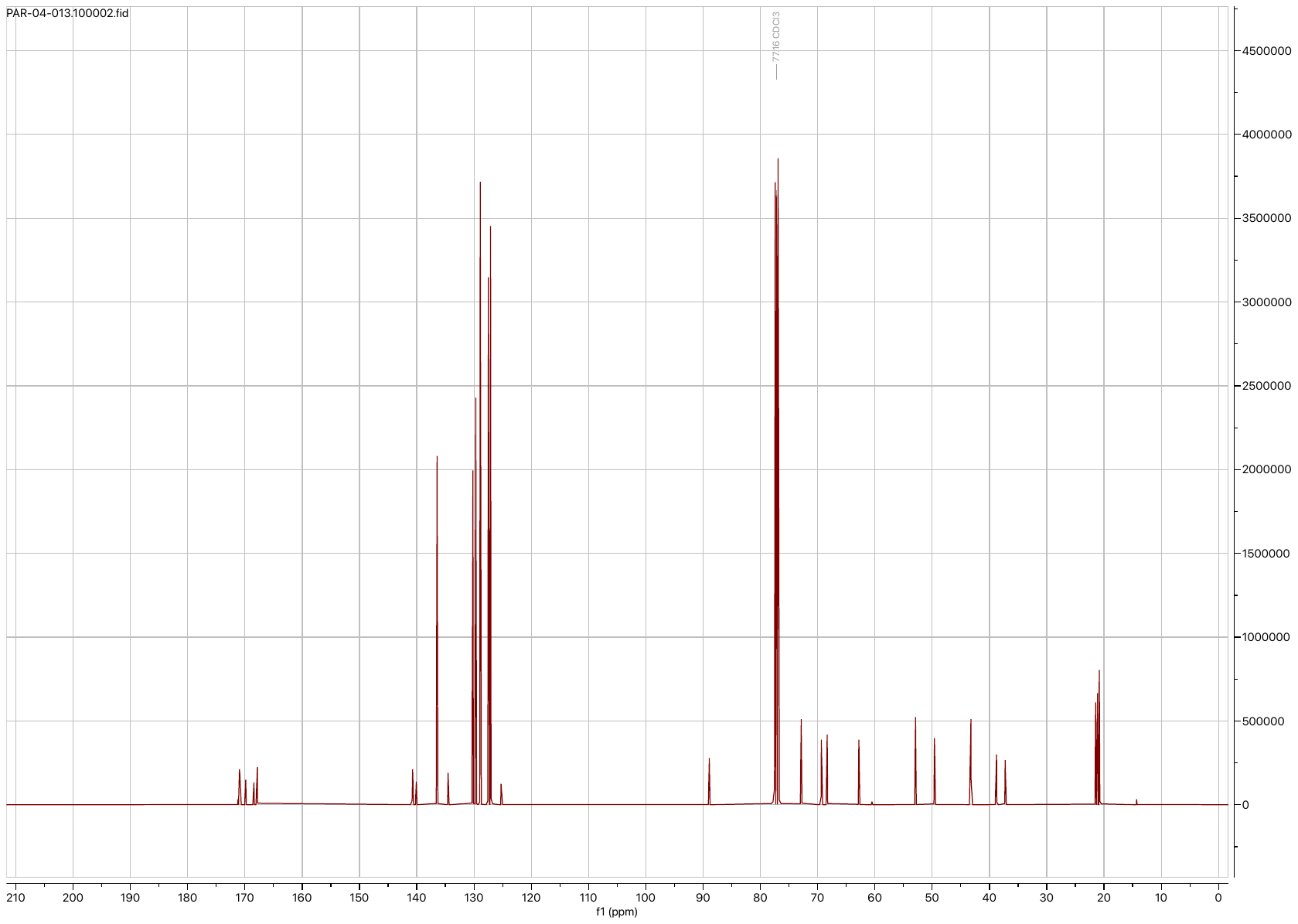

**S5 ^13^C NMR**

**S5 ^1^H NMR**

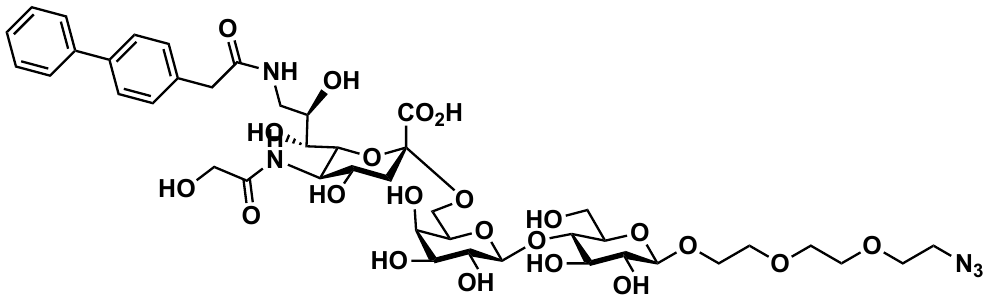

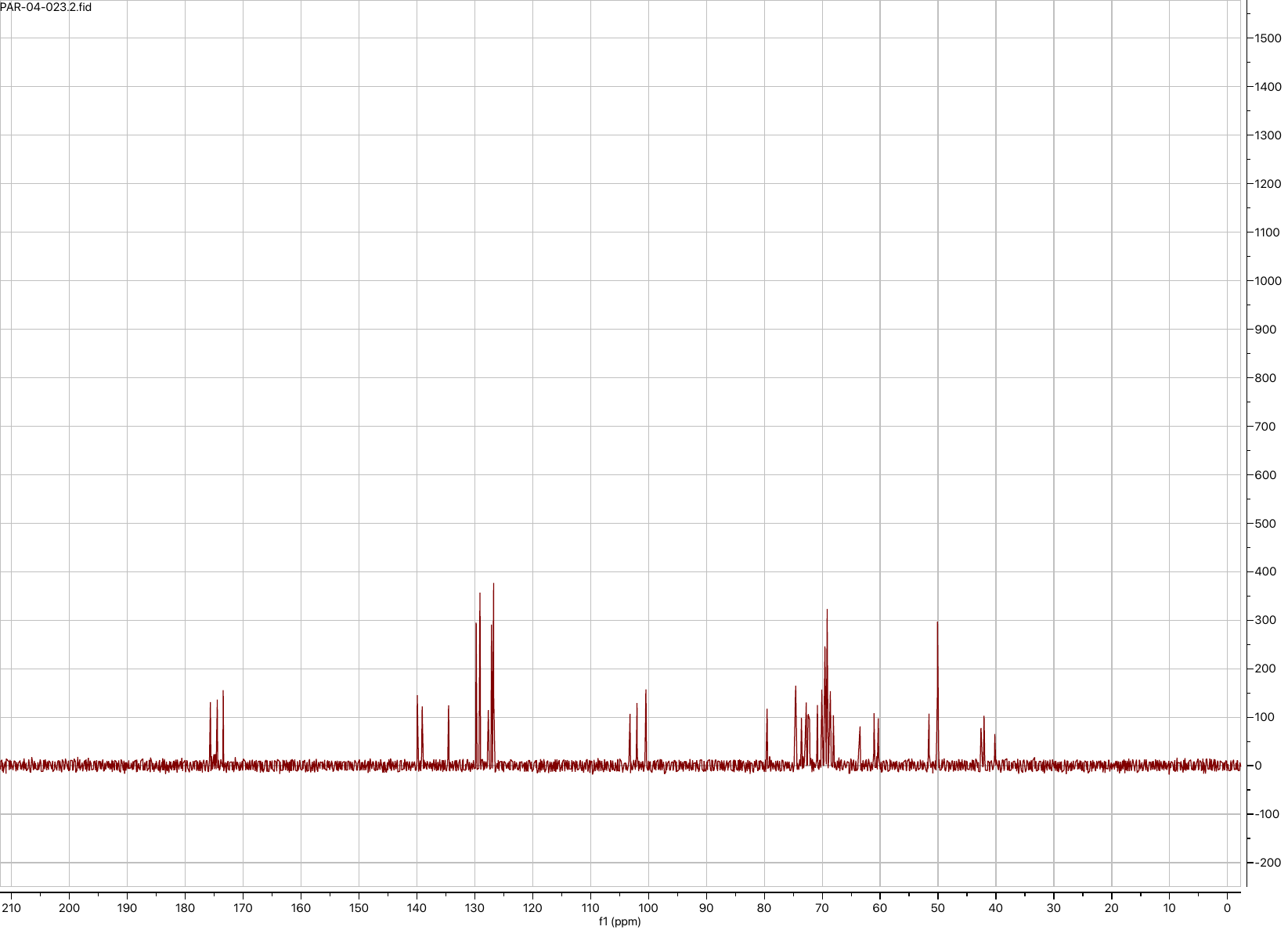

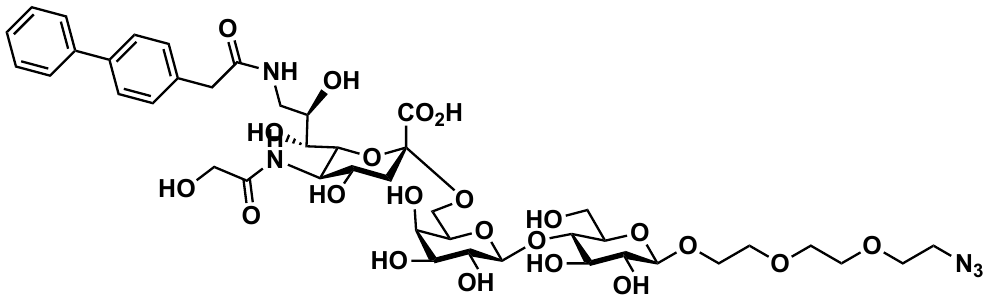

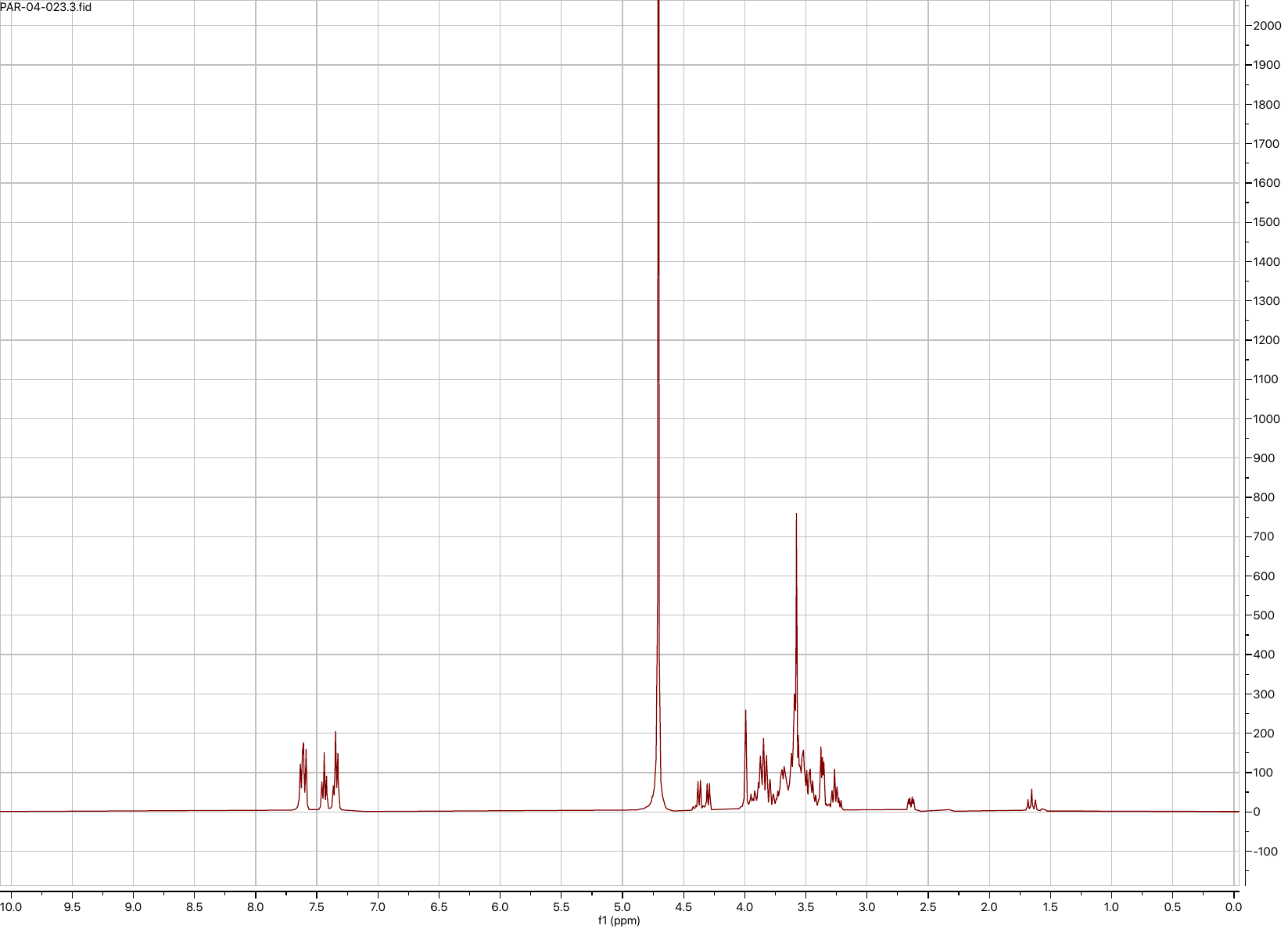

**mCD22L-azide ^13^C NMR**

**mCD22L-azide ^1^H NMR**
